# Wild emmer introgressions alter root-to-shoot growth dynamics in response to water stress

**DOI:** 10.1101/2020.06.17.157461

**Authors:** Harel Bacher, Feiyu Zhu, Tian Gao, Kan Liu, Balpreet K Dhatt, Tala Awada, Chi Zhang, Assaf Distelfeld, Hongfeng Yu, Zvi Peleg, Harkamal Walia

**Author notes:** Corresponding authors: Harkamal Walia, and Zvi Peleg.

## Abstract

Water deficit is a major limiting factor for wheat (*Triticum* sp.) development and productivity. One approach to increase water stress adaptation in wheat is incorporating novel alleles from the drought-adapted wheat progenitor, wild emmer (*T. turgidum* ssp. *dicoccoides*). We explored this idea in the context of vegetative growth by examining the phenotypic consequence of a series of wild emmer (*acc*. Zavitan) introgressions into elite durum wheat (*cv*. Svevo) under water-limited conditions. Using image-based phenotyping we cataloged divergent (from Svevo) growth responses to water stress ranging from high plasticity to high stability among the introgression lines. We identified an introgression line (IL20) that exhibits a highly plastic response to water stress by shifting its root-to-shoot biomass ratio for detailed characterization. By combining genotypic information with root transcriptome analysis, we propose several candidate genes (including a root-specific kinase) that can confer the shoot-to-root carbon resource allocation in IL20 under water stress. Discovery of high plasticity trait in IL20 in response to water stress highlights the potential of wild introgressions for enhancing stress adaptation via mechanisms that may be absent or rare in elite breeding material.

## INTRODUCTION

Wheat (*Triticum* sp.) is the most widely grown crop in the world, providing about one-fifth of the caloric intake by humans (FAOstat, 2017). To meet the food demand of the rising population, it is estimated that we will need at least a 60% increase in wheat production by 2050 (Myers et al., 2017). This is expected to be a major challenge, especially under projected climate change scenarios, and associated increases in variability of precipitation and frequency and intensity of drought in many agricultural regions (Cook et al., 2018). Genetic improvement in wheat, coupled with better agronomic management, is a core component for addressing this challenge. Key targets for enhanced adaptability are increased biomass accumulation during vegetative growth, enhanced water-use-efficiency and stability of yield parameters during reproductive growth under water-limiting environments (Araus et al., 2002). These can be realized at the molecular, morphological, and physiological level (Gupta et al., 2020). Increased genetic variability in breeding programs, particularly when the introduced variants can be associated with improved adaptation to water stress, can be a valuable resource for developing climate resilient wheat cultivars for the future.

From a physiological perspective improved adaptation to water-limited environments, especially with mild-to-moderate water stress, is inherently linked to higher biomass accumulation, which is a function of photosynthetic capacity at the canopy level. Canopy photosynthesis typically translates into higher yield in many crops and environments (Zelitch, 1982; Ashley and Boerma, 1989). Water stress results in a decline in turgor, cell division, and leaf growth, which also decreases the photosynthetic surface area and hence, overall photosynthetic capacity independent of the photosynthetic efficiency (Hsiao, 1973). Therefore, vegetative shoot growth in crops such as wheat can be considered an integrated trait on a temporal scale, with growth before reproductive transition impacting grain yields. Increasing genetic variation in wheat for rate of leaf area growth and duration of growth under water stress can be important yield determining trait under water stress (Richards, 2000). Capturing the phenotypic variation in shoot growth for genetic analysis requires accurate longitudinal measurements that are now feasible with non-destructive imaging platforms (Berger et al., 2010).

Relatively less is known about root responses to water stress due to inherently higher root plasticity and difficulty in accurately measuring their phenotypic traits. Wheat breeding is known to have reduced root size in modern varieties relative to wild ancestors or landraces in many environments (Waines and Ehdaie, 2007). This is likely due to higher photosynthetic cost of root growth and respiration that led to allelic enrichment that favours reduced carbon allocation to roots when selections are made under optimal environments (Lambers and Atkin, 1996). Under water stress, shoot growth is reduced as more carbon is allocated to roots, which results in higher root-to-shoot ratio (Correa et al., 2019). Deeper roots and more lateral root growth under water limited conditions enables plant access to more water during grain filling (Campos et al., 2004). The resulting greater stomatal conductance, cooler canopies and maintenance of physiological activity reduces grain yield losses in later developmental stages (Kirkegaard et al., 2007). Optimal root-to-shoot partitioning enables the balance between productivity and root water absorption (Voss-Fels et al., 2018). While the impact of the root-to-shoot allocation on drought tolerance and yield is likely to be context dependent, phenotypic plasticity in resource partitioning is an important trait to characterize from a genetic perspective for overall germplasm enhancement and may compensate for relatively lower allelic enrichment due to breeding selections made under optimal water conditions. The impact of root-to-shoot plasticity on grain yield will also depend on the seasonal precipitation profile and soil type among other factors.

One of the challenges with capturing the genetic variation underlying the plasticity in shoot and root growth in response to water stress is the temporal resolution needed for phenotyping a large number of accessions, making this intractable through manual, destructive measurements. High-throughput, image-based platforms can greatly improve the temporal resolution of phenotyping shoot responses to water stress across populations, although similar approaches for directly measuring roots responses are still limiting (Yang et al., 2020). Our ability to identify novel phenotypic responses and their genetic basis depends on the level of detectable phenotypic variation in the population and access to genomic resources. The range of phenotypic variation within a background or population can be increased significantly by introducing alleles from wild or related species [e.g. tomato (*Solanum lycopersicum*; Arms et al., 2015), barley (*Hordeum vulgare*; Baum et al., 2003) and rice (*Oryza sativa*; Tsujimura et al., 2019)]. Wild relatives, especially those adapted to semi-arid environments can be source of novel water stress responsive phenotypes that may be missing or rare in breeding germplasm.

In this study, we focused on wild emmer wheat [*T. turgidum* ssp. *dicoccoides* (Körn.) Thell.], the direct allotetraploid (2n = 4x = 28; genome BBAA) progenitor of all domesticated wheats (McFadden and Sears, 1946). Wild emmer thrives across the Near Eastern Fertile Crescent in a wide eco-geographic amplitude and harbors a rich allelic repertoire for numerous agronomic traits, including drought tolerance (Peleg et al., 2005). Introgression of wild emmer alleles has been shown to impact wheat adaptation to water stress (Golan et al., 2018; Merchuk-Ovnat et al., 2017). We have used a set of wild emmer introgression lines (ILs) in an elite tetraploid wheat background to discover novel phenotypic responses to water stress, using high temporal resolution imaging platform. We tested the hypothesis that introducing introgressions from wild emmer into durum wheat will expand the range of phenotypic responses to water stress with minimal loss in desirable agronomic traits of the elite cultivar. We identified two contrasting water stress response strategies, one where the phenotypic stability under water stress is observed and another that involves rapid change in carbon allocation relative to the elite cultivar. We characterized representative ILs for these two strategies for gaining further physiological insights. One of the ILs exhibiting a change in root-to-shoot ratio in response to water-stress was used for genetic and transcriptomic analysis to identify candidate genes localizing to the wild emmer introgressions. Our study highlights the potential of wild introgressions to promote various water stress responsive dynamics, as well as characterization of water stress adaptive mechanisms that can enhance climate resilience in wheat.

## RESULTS

### Wild emmer introgressions confer divergent water stress responses

The goal of this study was to investigate if the introduction of small wild emmer introgressions can expand the range of phenotypic response to water stress in an elite wheat cultivar background. To accomplish this, we developed a set of introgression lines (ILs) using elite durum wheat cultivar, Svevo as the backcross parent, and Zavitan as the source of wild emmer introgressions (Avni et al., 2014). Zavitan is well-adapted to the semi-arid environment and has a sequenced and annotated genome thus making it more accessible for downstream genetic analysis compared to other wild emmer accessions (Avni et al., 2017). The subset of ILs selected for this study by eliminating the wild alleles for the *Reduced height* (*Rht*)*-B1b* gene and non-brittle spike (*TtBtr1*) genes. This set included 47 wild emmer ILs were genotyped using the 90K SNP chip, and poses 1.3-14.2% of the Zavitan genes per IL (Oren, 2020). To examine their phenotypic responses to water stress using an automated high-throughput, image-based phenotyping platform. These ILs were grown under well-watered (WW; 80% field capacity) and water-limited (WL; 30% field capacity). Starting 11 days after transplanting (DAT) for 35 days of imaging, we collected five side-view images for each plant on each imaging day, to calculate the projected shoot area (PSA) from pixel counts and estimate daily shoot biomass accumulation as described in Campbell et al. (2015) and Knecht et al. (2016). The temporal PSA is a reliable proxy for shoot growth dynamics of the ILs and Svevo (Supplemental Fig. S1). We did not include Zavitan in this imaging experiment as it exhibits a weak growth habit when grown in pots under greenhouse conditions (observed during multiple seed increases). Since retention of desirable agronomic traits was a key criterion for initial selection of ILs for this study, we deemed the elite parent, Svevo to be a more suitable control for comparing phenotypic responses under WL treatment. Our observation that the PSA-based growth curves for Svevo were similar to the median response of all ILs collectively during the experiment supports this rationale. This suggests that the ILs biomass accumulation (derived from PSA) segregated around Svevo performance (Supplemental Fig. S2A). Other morphological traits such as plant width and convex area (architecture) also exhibited a pattern similar to PSA (Supplemental Fig. S2B-E). Although, most of the ILs reached the target of 30% field capacity in the WL treatment after an average of 19 DAT (ranged from 14 to 24 DAT) (Supplemental Fig. S3), many exhibited significant differences in biomass accumulation as early as 10 DAT (collectively), indicating an early water stress response as well as divergence in shoot growth response among the ILs (Supplemental Fig. S4).

To examine the consequences of early water stress response, we plotted the density distribution of the ILs under WW and WL treatments for several morphological traits at 35 DAT. The ILs exhibited a broad range for all the traits with Svevo positioned close to the average value for most traits (Fig. 1A). Overall, the ILs showed a strong reduction in PSA as evident from the separation of the distribution curves in response to water stress. A relatively lower separation between the WW and WL treatments was observed for additional morphological traits, such as plant width, WUE and density and. WUE was derived from daily increments in PSA and daily water use by the genotypes. The phenotypic distribution of plant height among the ILs under WL treatment varied more around the mean compared to the WW treatment. This was in contrast to the change in phenotypic distribution of PSA for the IL set, which varied less and was narrower under WL treatment. This could be due to plant height being one of the selection criteria for the ILs. Notably, the phenotypic range of WUE varied more under WL compared to the WW conditions.

**Figure 1.**
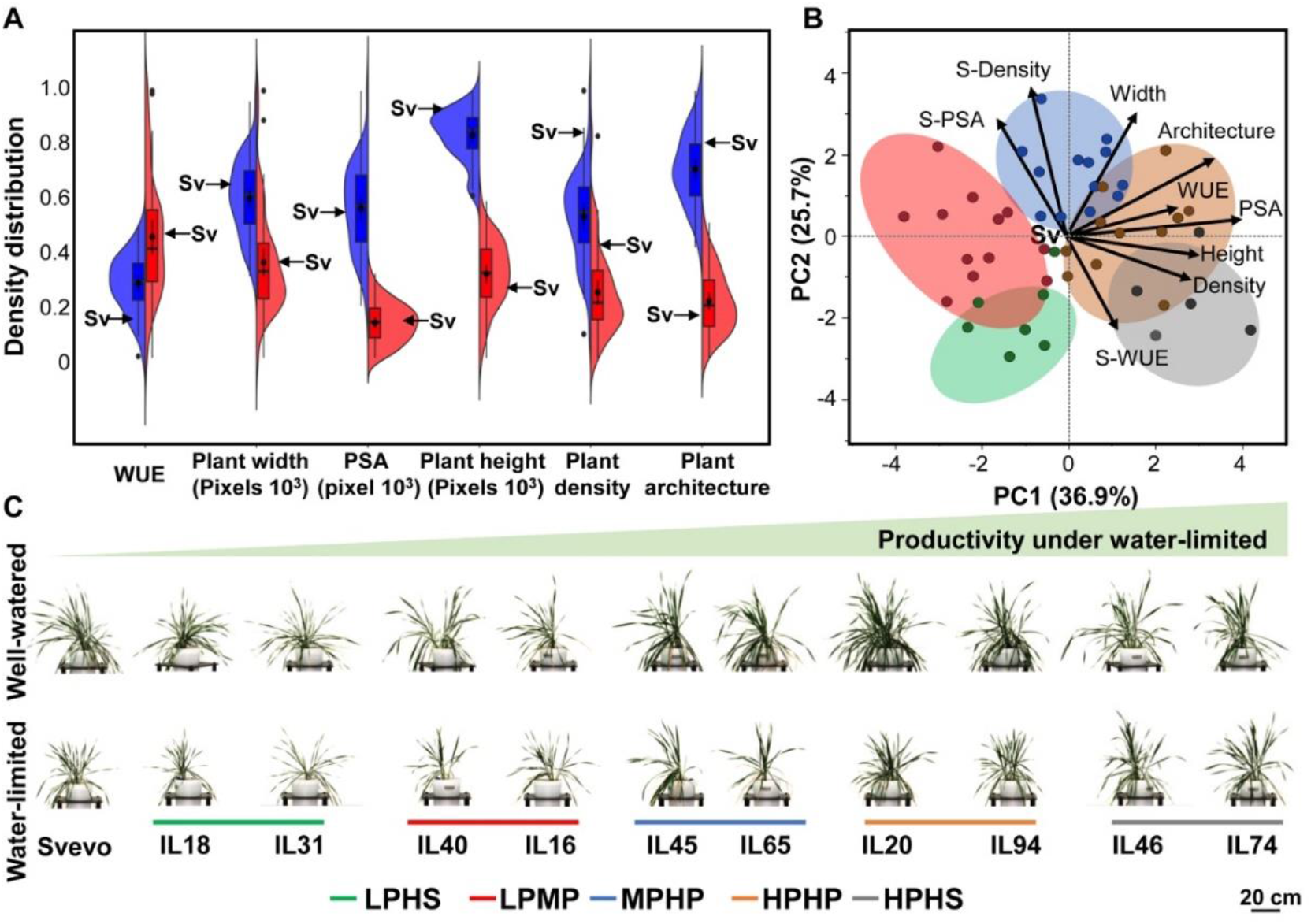
Wild introgressions promote phenotypic diversity. (**A**) Density distribution of morpho-physiological traits for 47 introgression lines (ILs) under well-watered (WW, blue) and water-limited (WL, red). Water-use efficiency (WUE), plant width, projected shoot area (PSA), plant height, plant density and plant architecture (convex area). The parental line Svevo (Sv) is marked with arrow. (**B**) Principal component (PC) analysis of morpho-physiological traits under WL conditions and expressed as drought susceptibility index (S). Biplot vectors are trait factor loadings for PC1 and PC2. The five clusters of stress responsiveness: high productivity - high stability (HPHS; gray), high productivity - high plasticity (HPHP; Orange), moderate productivity-high plasticity (MPHP; Blue), low productivity-moderate plasticity (LPMP; Red), low productivity-high stability (LPHS; Green). (**C**) Representative images of ILs from each responsiveness cluster under WW and WL treatments at 35 d after transplanting.

Understanding the relationship between the morpho-physiological traits can provide better insights into the key determinants of expanded phenotypic range observed among the ILs. Therefore, we performed correlation analysis between these traits at 35 DAT (Supplemental Fig. S5 and Supplemental Table S1). PSA was positively correlated with all morphological traits suggesting that plant biomass and architecture are tightly associated regardless of water availability. Under WL conditions, PSA and plant density were positively correlated with WUE, suggesting that plant architecture can affect the WUE under stress. Water stress was relatively more consequential in altering the inverse correlation between height and width, density, and WUE relative to WW treatment (Supplemental Fig. S5B). To further understand the response of plant phenotypic traits to WL treatment, we performed principal component analysis (PCA) of the morpho-physiological traits under WL treatment as well as in relative terms S (i.e., drought susceptibility index) (Fig. 1B). We define stress indices as a plant’s ability to maintain similar behavior under WL relative to its WW values for a given trait. PCA identified three major PCs (Eigenvalues > 1.2) accounting collectively for 76% of the phenotypic variance among the ILs (Supplemental Fig. S6). PC1 explained 36.9% of total variation and related positively with PSA, plant height, plant architecture, WUE, and plant density. PC2 explained 25.7% of the total variation and related positively with plant width, S-PSA, and S-density and negatively with S-WUE. PC3 explained 13.4% of the total variation and was positively related with WUE, S-PSA, and plant density. These results suggest that higher biomass and greater plant density were positively correlated with higher WUE under WL conditions. From the S-index perspective, high S-PSA which reflects significant biomass reduction due to water stress correlated with low S-WUE and confirmed that without WUE adaptation, plants will reduce their biomass gain under water stress.

Next, we sought to categorize the observed phenotypic divergence among the ILs with the goal to obtain more genotypic specificity associated with the phenotypic patterns (Fig. 1C). For this we performed hierarchical clustering analysis of the morpho-physiological traits under WL treatment and derived stress index traits (Supplemental Fig. S7). Clustering analysis separated the ILs into five distinct clusters, which we are broadly describe as following: Cluster 1 (high productivity and high stability, HPHS), Cluster 2 (high productivity and high plasticity, HPHP), Cluster 3 (moderate productivity and high plasticity, MPHP), Cluster 4 (low productivity and moderate plasticity, LPMP), and Cluster 5 (low productivity and high stability, LPHS). The productivity in the context of this analysis implies biomass accumulation under WL and plasticity is defined as the genotype’s ability to exhibit a relatively rapid change in a phenotypic trait in response to water stress. Svevo resolved to Cluster 4 (LPMP), which is characterized by low PSA and WUE, and an intermediate response to water stress. The two most productive clusters (HP), Cluster 1 and Cluster 2 showed differential stress response as expressed in the drought susceptibility index. Cluster 1 exhibited low S-PSA values indicative of a smaller change between WW and WL treatments. Cluster 2 has the highest WUE under WL and relatively high values of S-PSA, resulting in high plasticity for these ILs. Several Cluster 3 ILs have relatively higher value for convex area and plant width without a compensating decline in height. This increase in width and/or convex area without a compensating decline in plant height is also evident for two ILs in Cluster 1. Our clustering analysis enabled us to resolve ILs into various phenotypic categories and also identify representative ILs for each of the cluster for detailed characterization.

### Water stress responsiveness classification based on temporal growth dynamics

Although the clustering analysis using endpoint measurements of the ILs provides a useful perspective, the temporal dynamics for these traits that precede these phenotypic outcomes can enhance our understanding of water stress responses. For this, we mapped the overall trajectories and phenotypic distributions of these traits on a weekly basis (Fig. 2A). In general, all clusters exhibited higher biomass accumulation and higher coefficient of variance (CV) under WW relative to WL treatment (Fig. 2A; Supplemental Table S2). The PSA distributions under WW and WL treatments showed that the high stability (HS) clusters exhibited substantial overlap between the WW and WL curves in weeks 5 and 6. The point of significant response to water stress was determined when three contiguous days of significant (*P*≤0.05) difference in growth between treatments was recorded, which ranged between 10 DAT (HPHP cluster) to 26 DAT (HPHS cluster) (Fig. 2A; Supplemental Table S3). A similar pattern was found for plant architecture and density (Supplemental Fig. S8). The parental line (Svevo; LPMP cluster) expressed an intermediate response (17, 18, and 15 DAT for PSA, plant architecture, and density, respectively; Supplemental Fig. S8). Although the MPHP cluster exhibits high biomass accumulation under WW treatment, it was labeled as moderate productivity (MP) based on its performance under WL treatment. The cluster’s classification to productivity (i.e., HP, MP, and LP) was found to be significantly different (*P*<10^−4^) under WL. This analysis enabled us to capture the temporal dynamics, which are typically challenging to determine without extensive destructive sampling but important to examine the level of plasticity under varying environmental conditions.

**Figure 2.**
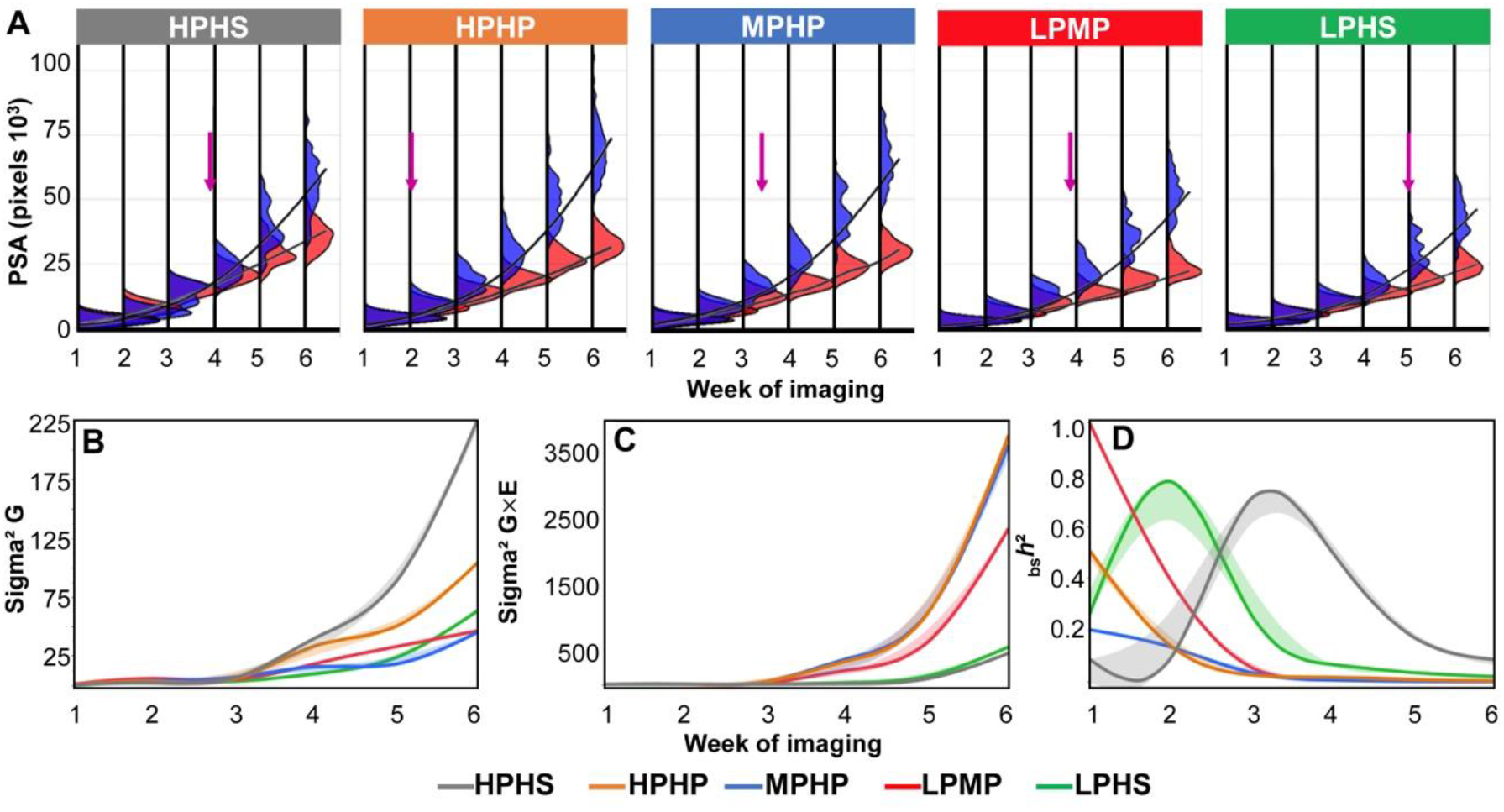
Longitudinal dynamics of the five responsiveness clusters. (**A**) Longitudinal frequency distribution of biomass accumulation (projected shoot area; PSA) of each water stress responsive cluster under well-watered (WW; blue) and water-limited (WL; red) treatments. The five clusters are: high productivity-high stability (HPHS; gray), high productivity-high plasticity (HPHP; Orange), moderate productivity-high plasticity (MPHP; Blue), low productivity-moderate plasticity (LPMP; Red), low productivity-high stability (LPHS; Green). The time point of significant difference in response to water stress is marked with an arrow (*P*≤0.05). Longitudinal heritability components of (**B**) genetic (Sigma^2^ G), (**C**) environmental interaction (Sigma^2^G × E), and (**D**) broad-sense heritability (_*bs*_*h*^2^). Continues line represent the smooth curve through the data and the shaded area represents the standard error of the smooth curve.

### Plant responsiveness clusters expressed in heritability dynamics

To dissect the genetic (G) and environmental (E) components of PSA underlying each responsiveness cluster through studied developmental stages, we calculated broad-sense heritability and its components. The HPHS cluster exhibited the highest genetic component (sigma^2^ G), which increased with the progression of water stress duration (Fig. 2B). On the other hand, HPHP and MPHP clusters had lower genetic components and the highest G×E interaction (Sigma^2^ G×E) (Fig. 2B, C). The broad-sense heritability dynamics (_*bs*_*h*^2^) of PSA showed clear separation into stability (LPHS and HPHS) and plasticity (LPMP, MPHP, and HPHP) (Fig. 2D). In general, the level of PSA _*bs*_*h*^2^ decreased over time. Heritability dynamics of plant density showed a strong genetic component for HPHP and a high environmental effect for LPMP that increased over time. Plant architecture presented a high environmental effect on MPHP, causing low _*bs*_*h*^2^ for this cluster (Supplemental Fig. S9). Overall, the heritability dynamics of the responsiveness cluster emphasized that stability and plasticity derived from both genetic and environmental effects within the IL panel.

### IL20 exhibited higher assimilation rate under water-limited conditions

We next focused on the phenotypic plasticity/stability under WL, by comparing two high productivity clusters HPHP and HPHS, represented by IL20 and IL46, respectively, for downstream physiological analysis. We targeted the temporal window during the early growth stages (15-19 Zadocks scale; Zadocks et al., 1974), and used the same experimental design that previously enabled us to categorize ILs based on growth rate and water stress response. Under WL treatment, the relative growth rate dynamics demonstrated the advantage of the two productive clusters as expressed in higher linear equation slope 452.3 for IL20 and 558.1 for IL46, compared to Svevo (284.45; *P*<0.005) (Fig. 3A; Supplemental Table S4). Under WW treatment, only IL20 had higher slope compared to Svevo (*P*=0.001). While IL46 maintained a similar linear equation slope under both water treatments (representing the high stability cluster), IL20 exhibited a significant change in the regression pattern, from 849.75 under WW to 452.32 under WL (*P*<10^−3^) (Fig. 3A) consistent with its selection based on high plasticity.

**Figure 3.**
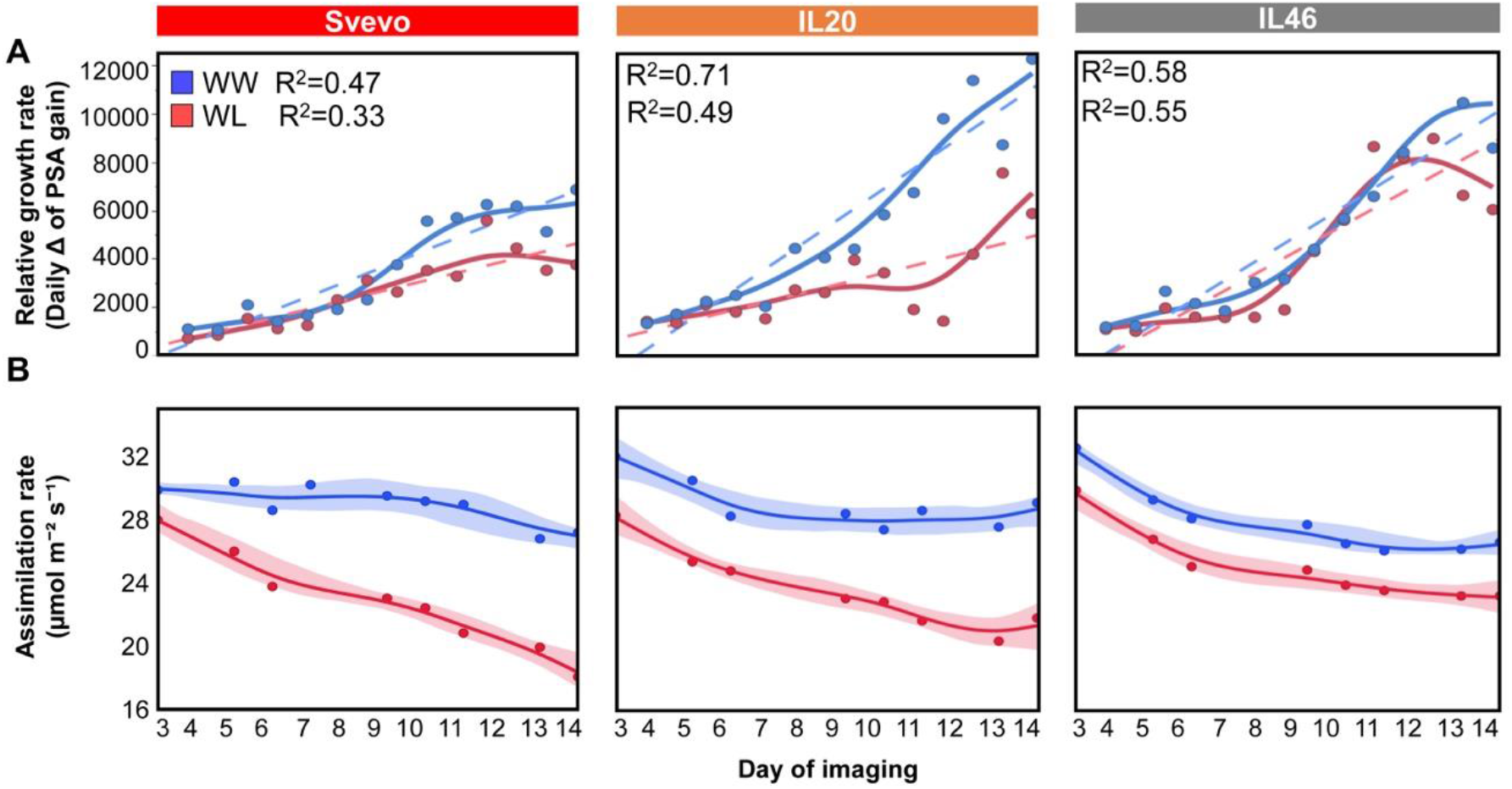
Longitudinal dynamics of Svevo, IL20 and IL46 for (**A**) relative growth rate, (**B**) net assimilation rate under well-watered (WW; blue) and water-limited (WL; red) treatments. Dashed lines represent the fitted linear growth of each genotype under specific water treatment. Markers represents the genotypic mean under specific water treatment (*n*=4). Continuous line represents the smooth curve through the data and the shaded area represents the standard error of the smooth curve.

To complement the imaging of the two ILs and Svevo, we also measured gas exchange parameters over eight time points during the course of the experiment. Average assimilation rate (A) declined with the progression of water stress as expected, with Svevo exhibiting the most reduction (37.7%), whereas the high stability IL46 had only 22.9% reduction (Fig. 3B). Notably, IL20 exhibited the highest assimilation rate under WW treatment over time (30.19 μmol m^−2^ s^−1^), whereas under WL both IL46 and IL20 exhibited similar A (25.35 and 24.26 μmol m^−2^ s^−1^, respectively), that was significantly higher than Svevo (22.66 μmol m^−2^ s^−1^; *P*<0.047; Supplemental Fig. S10). IL20 also maintained significantly higher stomatal conductance (*g*_*s*_) (*P*=0.013) and transpiration rate (E) (*P*=0.024) under WL relative to Svevo (Supplemental Fig. S11, Supplemental Table S5). Under WW treatment, both IL20 and IL46 had higher (*g*_*s*_) compared to Svevo.

### IL20 exhibits higher root-to-shoot ratio under water stress

Since IL20 maintained higher assimilation rate and stomatal conductance under WL treatment, we investigated if this was related to improved water uptake due to differential root growth response under water stress (Fig. 4A). We measured root dry weights from 21 DAT plants grown in potted soil and found that both IL46 and IL20 had higher root biomass relative to Svevo (*P≤*0.001) under WW treatment. However, under WL treatment, Svevo root biomass were significantly lower than IL20 (*P*=0.003). Further, IL20 also exhibited a higher root-to-shoot ratio when compared with Svevo under WL conditions (*P*=0.046) (Fig. 4B, C; Supplemental Table S6). Our data suggests that the root response of IL20 under water stress diverges from Svevo. To explore this differential root response to water stress in a developmental context, we performed a seedling stage assay using a paper roll set-up (Placido et al., 2013). While the shoot length of IL20 and Svevo was similar under WW and WL treatments, IL20 exhibited significantly higher root length throughout the experiment, with 10.3% longer roots at the end of the experiment (25.21 *vs.* 22.85 cm, for IL20 and Svevo, respectively; *P*=0.006) under WL. This advantage expressed in the higher (12.5%) root-to-shoot ratio of IL20 compared with Svevo on the last day (*P*=0.001; Supplemental Fig. S12A-F). This suggests that the root growth dynamic of IL20 is different from Svevo even during early seedling stage and more apparent under WL treatment and results in increase in the root-to-shoot ratio. These results show that root biomass in later stages (19 in Zadocks scale) and root length at the seedling stage (11 in Zadocks scale) have a similar response in IL20 under WL treatment (Fig. 4, Supplemental Fig. S12).

**Figure 4.**
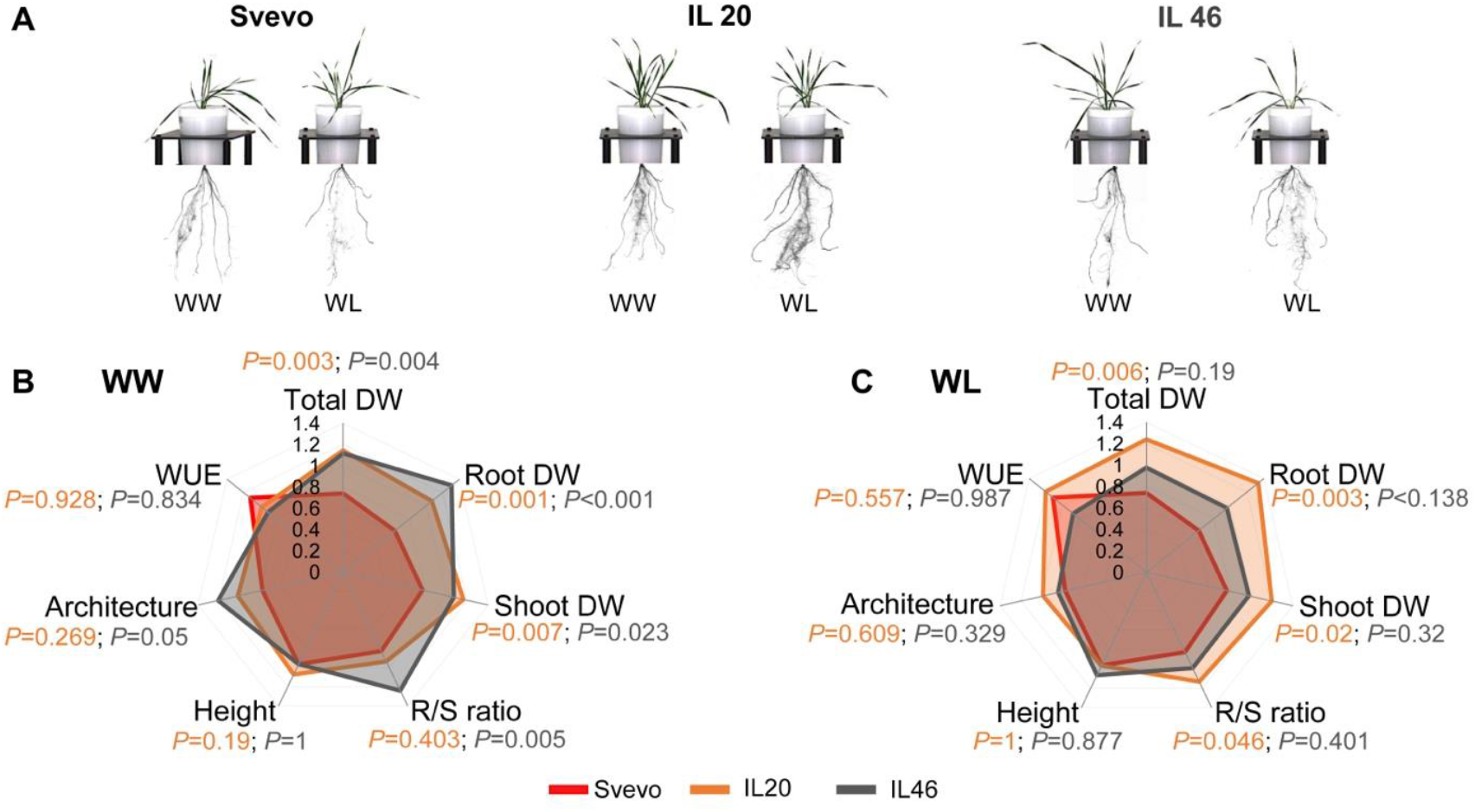
Morpho-physiological modification in response to water stress. (**A**) Representative image of Svevo, IL20, and IL46 under well-watered (WW) and water-limited (WL) treatments 14 d after transplanting. Radar charts comparing the phenotypic traits of Svevo (red), IL20 (orange), and IL46 (gray) plants under (**B**) WW and (**C**) WL treatments. Values are means (*n*=4). Total dry weight (Total DW), water-use efficiency (WUE), plant architecture (convex area), plant height (height), root-to-shoot ratio (R/S ratio), shoot DW, and root DW.

### Transcriptome analysis of IL20 and Svevo roots in response to water stress

Given the differential root growth and the root-to-shoot ratio between Svevo and IL20 in the seedling stage, we reasoned that the underlying gene(s) responsible for these phenotypes could be the same, resulting in similar root-to-shoot ratio plasticity that was observed in later vegetative stages. Therefore, we performed a transcriptome analysis on roots from the seedling stage experiment to identify candidate genes that underlie the root-to-shoot plasticity phenotype. Seedling root sampling for transcriptome is more precise as it limits gene transcript changes caused by root damage that occurs with sampling roots from older plants growing in soil or sand. We combined the transcriptome analysis with the genotypic data of IL20 and Svevo to map the differentially expressed genes (DEGs) to specific introgressions. IL20 has six introgressions from Zavitan, the wild emmer parent, distributed on five chromosomes (Table S7), accounting for ~4.5% from the Zavitan genome (Avni e al., 2017). Based on public annotations, a total of 651 genes from the homozygous regions (Supplemental Table S8) map to these introgressions. Under WL treatment, when the root phenotype is most apparent, we identified 599 DEGs (Fig. 5A) between Svevo and IL20, with 37 genes (6.17%) co-localizing to the introgressions. Of these, 425 genes were down-regulated and 174 genes were up-regulated in IL20 (Supplementary Table S9). Under WL treatment, 39.23% of the DEGs were differently expressed between Svevo and IL20 (56 up- and 179 down-regulated), whereas only 11.35% were expressed differently under WW treatment.

**Figure 5.**
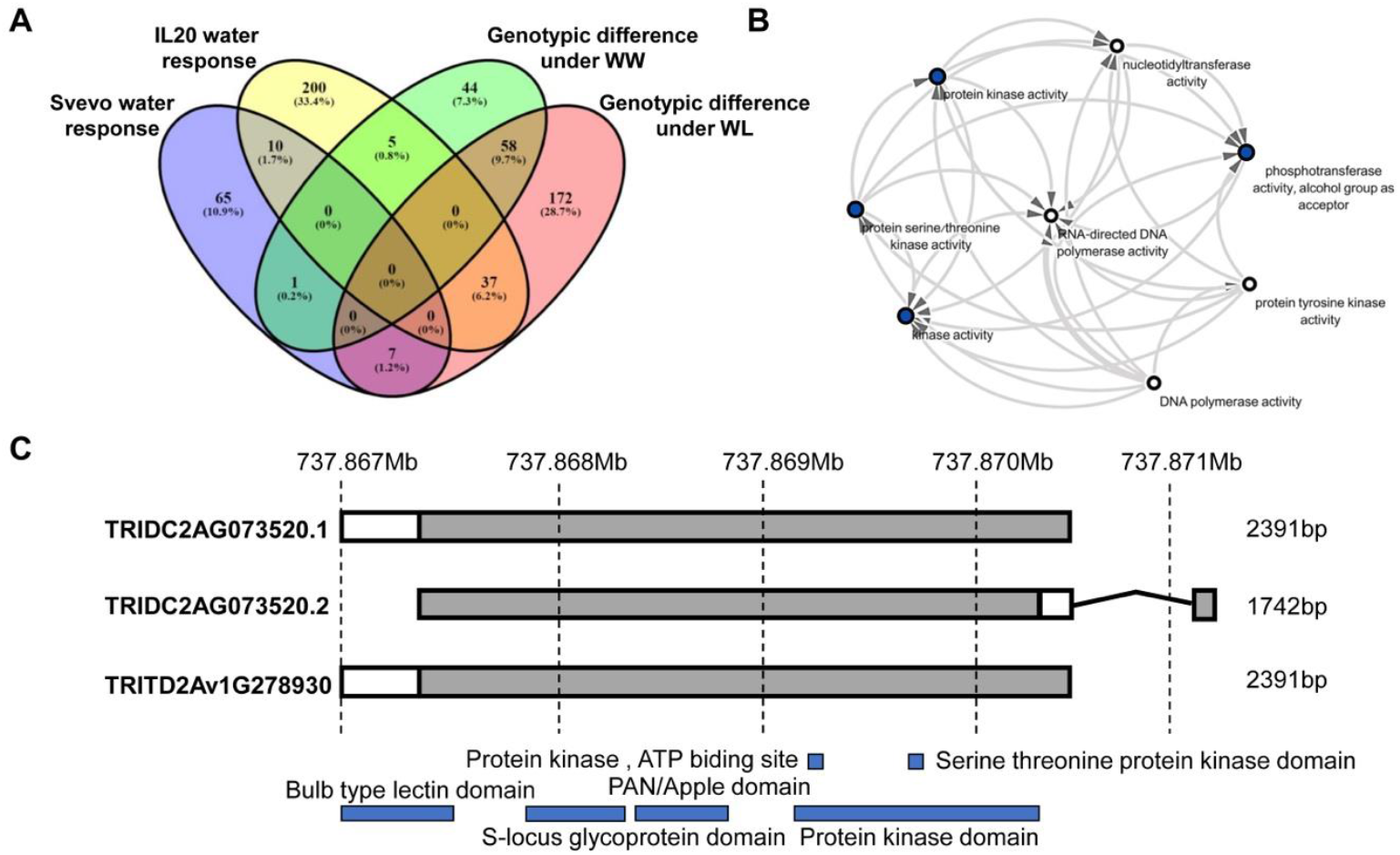
Differently expressed genes (DEGs) comparison for IL20 and Svevo. (**A**) A four-way Venn diagram of DEGs among IL20 and Svevo under well-watered (WW) and water-limited (WL) treatments. (**B**) Network co-expression pattern of the DEGs associated with kinase activity. Node with blue color represents higher interaction. (**C**) Splice variation of TRIDC2AG073520 gene, a candidate for root phenotypes of IL20 that maps to the introgression. TRITD2Av1G278930 represent Svevo allele for the current gene.

### Candidate genes associated with longer roots under water stress

We next examined the differentially abundant transcript(s) that localize to the introgressions in IL20 and identified 17 DEGs under WW and 18 DEGs under WL treatments between IL20 and Svevo. Two DEGs (TRIDC4AG049220 and TRIDC4AG049940) were found to express only in IL20 in response to water stress. Given the IL20 root phenotype, we targeted root-related DEGs from this set, which yielded five candidate genes (CG; Supplementary Table S10). The criteria used to filter these five genes are based on literature searches of orthologs with root-associated phenotypes. Three of these genes were up-regulated in IL20 under WL (TRIDC4AG046080, TRIDC4AG048600, and TRIDC2AG073520), one gene was down-regulated under WL (TRIDC4AG046660) and one gene (TRIDC4AG046110) showed up-regulation under WW treatment only. Of these five genes, TRIDC4AG046080 is a low confidence gene based on the annotation of the Zavitan genome. The remaining four genes either have a SNP (TRIDC2AG073520, TRIDC4AG048600), or carry multiple polymorphisms (TRIDC4AG046660), or were presence/absence variation between the Zavitan and Svevo genomes (TRIDC4AG046110) (Supplementary Table S10). TRIDC4AG046110 encodes a *FAR1-RELATED SEQUENCE 4-like* isoform that is down-regulated in salt-susceptible sweet sorghum (*Sorghum bicolor*) roots (Yang et al., 2018). TRIDC4AG048600 is a *SIMILAR TO RCD ONE 1* (*SRO1*) gene. In *Arabidopsis* (*Arabidopsis thaliana*), a double mutant of *AtSRO1* exhibited shorter roots and a smaller cell division zone compared to wildtype plants (Teotia and Lamb, 2011). Sequence alignment of this gene against the Zavitan genome indicates a truncated protein in the Zavitan genome that may result in loss of function or a modified function.

The remaining three DEGs were associated with protein kinase function (Supplementary Table S10), where network analysis of molecular functions showed significant downstream transferase activity elements in various kinase activities (Fig. 5B). TRIDC4AG046080 is a homolog of a rice domain of the unknown function (*DUF581*) that, in *Arabidopsis*, was found to play a role in sucrose non-fermenting-related kinase (*SnRK1*) (Nietzsche et al., 2016). TRIDC4AG046660 is a Leucine-rich repeat receptor protein kinase (LRR-RLK) and TRIDC2AG073520 is a G-type lectin S-receptor-like serine/threonine-protein kinase (RLK). We examined the sequence of TRIDC2AG073520 in the Zavitan genome (Avni et al., 2017) and identified two splice variants on chromosome 2A, which are 2391bp and 1742bp for TRIDC2AG073520.1 and TRIDC2AG073520.2, respectively. In contrast, only a single variant (2391bp) was found in the tetraploid durum wheat (*cv*. Svevo) (Fig. 5C) as well as among 10 hexaploid bread wheat cultivars genomes (Appels et al., 2018; Walkowiak et al.,2020). This CG was mapped in the expression atlas of the Zavitan tissue-specific gene and suggest that it could be the primary candidate gene associated with the root-to-shoot ratio difference exhibited by IL20 (Supplemental Fig. S13).

## DISCUSSION

Wild plants have developed various reversible and non-reversible phenotypic plasticity strategies to cope with environmental uncertainty. Selection by humans, often under less variable environmental conditions has likely resulted in higher crop-plant phenotypic stability (Lopes et al., 2015). Consequently, many modern cultivars may have lost some of the fitness components needed for adapting to climate driven variation in many regions (Kissoudis et al., 2016). Wild ancestors of modern crops offer a promising source for genetic diversity and novel drought adaptive traits (Peleg et al., 2005; Golan et al., 2018).

The introgression of Zavitan alleles into a modern durum cultivar promoted higher phenotypic diversity under both WW and WL treatments, as expressed in plant architecture and biomass accumulation (Fig. 1). Although the IL panel was developed from a single wild emmer accession (Zavitan), yet it resulted in wide segregation of morpho-physiological traits (either positively or negatively). This accession originated from a habitat with high soil moisture fluctuations, due to a shallow brown basaltic soil type, which has been shown to promote diversity (Poot and Lambers, 2008; Peleg et al., 2008). This phenotypic variation is associated with the quantitative nature of these traits and the different combinations of wild and domesticated alleles. Interestingly, the mean biomass accumulation trajectory over time of the IL panel was similar to Svevo under both water treatments.

Water stress reduced biomass by around 50% (i.e., PSA) and altered plant architecture (i.e., convex area 12.5-48.5%) relative to the WW treatment (Supplementary Fig. S2), with both variables being positively associated with one another (Supplementary Fig. S5). Increased phenotypic variation in response to water stress was quantified by the calculation of drought susceptibility index (S-index). The combination of IL performance under WL with their S-indexes resulted in five distinct clusters of high phenotypic stability (HPHS, LPHS) and phenotypic plasticity (HPHP, MPHP, LPMP). Phenotypic stability is often associated with small changes in plant performance in response to unfavorable conditions. Escape (i.e., rapid growth to avoid the stress) is a common strategy of wild plants in xeric habitats and has been repeatedly reported for many wild grasses such as emmer wheat (Peleg et al., 2005), *Brachypodium distachyon* (Opanowicz et al., 2008), and *Avena barbata* (Sherrard and Maherali, 2006). Accordingly, the two clusters exhibiting phenotypic stability had biomass reductions of only 45 and 40% for LPHS and HPHS, respectively. Interestingly, the LPHS had characteristics of “small plants” (PSA, 50.4, and 27.8 kPixel for WW and WL, respectively), whereas HPHS had high biomass under WW and the highest values among all clusters under WL (67.1 and 40.6 kPixel, respectively). These results suggest that the phenotypic stability strategy is not size-dependent, but rather an active mechanism that enables plants to cope with water stress.

Wild emmer wheat populations were found to harbor rich phenotypic diversity for drought-adaptive traits, which correspond with the wide inter-annual and seasonal fluctuations in soil moisture availability of the Mediterranean basin (Peleg et al., 2005). Accordingly, the phenotypic plasticity clusters exhibited a greater reduction in biomass accumulation (55 and 56% for MPHP and HPHP, respectively). The HPHP cluster had the highest biomass under WW (PSA 81.8 kPixel); while under WL it exhibited more reduction, although biomass was still relatively high (36.4 kPixel) compared to all clusters.

Plant acclimation to water stress elicited physiological, morphological, and metabolic responses that occurred through coordinated spatio-temporal processes. These processes changed the physiological status of plants toward a new steady-state level that supports growth and fitness under unfavorable conditions. The temporal characterization of the responsiveness clusters showed that clusters with high phenotypic plasticity responded earlier (12, 8, and 10 kPixel for LPMP, MPHP, and HPHP, respectively) than those in high stability clusters (20 and 26 kPixel for LPHS and HPHS, respectively) (Fig. 2A). To understand the longitudinal genetic architecture of the responsiveness clusters, we calculated broad sense heritability (_*bs*_*h*^2^) dynamics. While the plasticity clusters exhibited a decrease in PSA _*bs*_*h*^2^ over time as a consequence of high G×E interaction (Sigma^2^ G×E) and low genetic component (Sigma^2^ G), the more phenotypically stable clusters showed increased heritability during early growth and decreased heritability at later stages, which corresponds to the delayed stress responses (Fig. 3).

Plants exhibit morphological and physiological adjustments to maintain water status and carbon assimilation under water stress (Chaves et al., 2009). The two high productivity clusters (HPHS and HPHP) exhibit contrasting response mechanisms, with the plasticity cluster responding earlier (16, 17, and 8 d early for PSA, plant density and plant architecture, respectively; Fig. 2; Supplementary Fig. S8). Detailed characterization of representative accessions for these two clusters (represented by IL20 and IL46 for HPHP and HPHS, respectively) confirmed the earlier response of HPHP in terms of relative growth rate (Fig. 3A), thus suggesting size-independent plant responsiveness to water stress. Consistent with the growth phenotype, IL46 maintained similar photosynthetic and transpiration rates under WW and WL, while IL20 responded as early as day 12, limiting its assimilation rate. Notably, IL20 had the highest photosynthetic rate under WW and exhibited a larger reduction under WL yet was able to maintain significantly higher assimilation rate than Svevo.

A fast stress responsiveness strategy may negatively affect carbon assimilation and growth; on the other hand, early acclimation can trigger a metabolic shift of carbon allocation to different plant organs (Rodrigues et al., 1993; Bohnert and Sheveleva, 1998). Thus, under water limitation, root-to-shoot ratio plasticity can mediate optimal resource partitioning between growth and development (Shipley and Meziane, 2002; Voss-Fels et al., 2017). Modern bread wheat cultivars have lower root-to-shoot ratios as compared to landraces (Siddique et al., 1990). Moreover, a comparison among wild emmer, domesticated emmer, and durum wheat showed a trend of reduced root-to-shoot ratio during the initial domestication from wild to domesticated emmer, and during wheat evolution under domestication (Gioia et al., 2015; Roucou et al., 2018). Accordingly, the introgression of alleles from Zavitan in the background of the elite durum wheat cultivar significantly increased the root-to-shoot ratio (30%) under WL as compared with the parental line (Fig. 4C). Likewise, Merchuk-Ovnat et al. (2017) reported a higher root-to-shoot ratio in response to water stress from wild emmer (*acc*. G18-16) introgression in the background of elite bread wheat cultivar. Thus, introducing new genetic diversity for root-to-shoot ratio plasticity from wild progenitors will facilitate the resilience of modern wheat cultivars to the projected fluctuating water availability during the growing season.

The root system is the site of interactions with the rhizosphere; thus, root architectural plasticity (i.e., allocational, morphological, anatomical, or developmental) is a critical adaptation strategy to environmental cues (Rellán-Álvarez et al., 2016; Golan et al., 2018). To better understand the genetic mechanism associated with the increased root biomass of IL20, we analyzed transcriptome response of roots under water stress. In general, differential transcriptional response of IL20 WL was greater than Svevo (223 *vs*. 73 DEGs, respectively). This is consistent with a previous study where miRNA expression in the roots of two wild emmer accessions (TR39477 and TTD-22) were significantly higher compared with domesticated durum wheat (*cv*. Kızıltan) under water stress (Akpinar et al., 2015). These results emphasize the potential of higher plasticity in transcriptional response of wild relatives compared to the domesticated germplasm.

Downstream gene network analysis highlighted the key role of protein kinases as hubs of interaction (Fig. 5B). Three CGs (TRIDC4AG046080, TRIDC2AG073520, and TRIDC4AG046660) were found associated with protein kinase function that mediates plant hormone and nutrient signaling, and cell cycle regulation (Laurie and Halford, 2001; Virlet et al., 2017). TRIDC4AG046660 is a leucine-rich repeat receptor-like protein kinase (LRR-RLK). Mutants of this gene in *Arabidopsis* (*At*2g33170) controls root growth and are mediated by cytokinin (Colette et al., 2011). TRIDC4AG046080 (DUF581 in rice) interacts with *SnRK1* and regulated by hormones and differentially regulated by hormones and environmental signals (Nietzsche et al., 2016). Wheat mutants containing a conserved DUF581 domain revealed a salt-induced gene (TaSRHP). Early stages of salt stress typically have an osmotic stress component that is similar to water stress. Over-expression of this gene in wild-type *Arabidopsis thaliana* resulted in enhanced resistance to both salt and drought stresses (Hou et al., 2013).

TRIDC2AG073520 (TRITD2Av1G27893 in Svevo) is a G-type lectin S-receptor-like serine/threonine-protein kinase gene. The domesticated allele contains a nonsynonymous mutation expressed as an amino acid shift (isoleucine to threonine). This CG was significantly up-regulated under WL in IL20 (FC 2.29, *P*_adj_=0.03). In *Arabidopsis*, drought and salinity stress-induced up-regulation of the gene (Sun et al., 2013). Moreover, the gene was expressed specifically in root tissue from the early seedling stage to 50% of ear emergence (Supplemental Fig. S14; Ramírez-González et al., 2018). Genetic dissection showed that the genomic region of this gene overlaps with a QTL affecting lateral root number per primary root (Maccaferri et al., 2016).

Two splice variants of TRITD2Av1G278930 were identified in the wild emmer genome (TRIDC2AG073520.1 and TRIDC2AG073520.2) and these included several polymorphisms in each variant. The TRIDC2AG073520.1 variant is similar to the domesticated variant, although it contains a nonsynonymous SNP. This gene was compared to the wheat pan-genome (Walkowiak et al., 2020) and similar SNP was found in all genomes compare to Zavitan, suggesting variation between wild and domesticated wheats. The TRIDC2AG073520.2 variant is different in length and exon number; however, the domains remain similar to the domesticated variant and the additional exon does not encode for a specific known domain (Fig. 5C). The underlying mechanism by which the identified splice variance and/or amino acid substitution affect wild emmer response to stress via longer root systems are yet to be discovered.

### Concluding remarks and future perspective

In this study, we show that targeting small and hence more genetically tractable wild introgressions can still yield surprisingly divergent phenptypic responses to water stress even after selection of ILs that have agronomically viable phenology. Our detailed physiological characterization combined with temporal phenomics apporaches provides novel insights into the divergent water stress response dynamics in an elite durum background. Although, we focused specifically on one IL for downstream characterization of its root-to-shoot plasticity response under water stress and candidate gene discovery, this work lays the foundation for characterization of additional lines from this IL panel and other introgressions in wheat. Collectively, our results suggest that incorporating the wild gene/alleles can enable greater phenotypic plasticity and has the potential to enhance environmental stress resilience.

## MATERIAL AND METHODS

### Plant material and experimental design

The wild emmer accession Zavitan was collected at the Zavitan nature reserve (32°56’24.52’’N, 35°42’10.56’’E; 288.4 m above sea level), Israel and was selected as the donor parent due to its robust morphology and drought tolerance (Avni et al., 2014). Uniform seeds of 47 wild emmer wheat (*acc*. Zavitan) introgression lines (IL) in the background of elite durum wheat (*cv*. Svevo) and their recurrent parent were used for the current study. A recombinant inbred line population from cross between durum wheat (*cv*. Svevo) and wild emmer (*acc*. Zavitan) was previously developed (Avni et al., 2014). Adapted RILs (i.e. with the genetic composition of post-domestication alleles of dwarfing gene *Reduce height* (*Rht*)*-B1b* and non-brittle spike genes (*TtBtr1-A and TtBtr1-B*), were selected and backcrossed three time and selfed over three generations. The 47 ILs were genotyped (Infinium iSelect 90K SNP chip array; Wang et al., 2014), resulting is equal SNP distribution across the genomes. Altogether, the ILs population cover 99.1% of the wild emmer Zavitan genome (Oren, 2020). Detailed information for the ILs panel is provided in Supplementary Table S11. Seeds were surface disinfected (1% sodium hypochloric acid for 30 minutes) and placed in Petri dishes on moist germination paper (Anchor Paper Co., St. Paul, MN, USA) about 3 cm apart, at 24°C in the dark for 5 days. Three uniform seedlings from each line were transplanted to a single pot (2L, 45×19.5 cm) filled with 1.2 kg of Fafard germination soil (Sungro, Massachusetts, USA), with osmocote fertilizer and Micromax micronutrients. Six days after transplanting (DAT), plants were thinned to one plant per pot. Pots were placed on automated carriers in the greenhouse (22/16°C day/night) and watered daily to 80% field capacity until the beginning of the experiment (11 DAT), 2-4 in Zadocks scale (Zadocks et al., 1974). The growth stages were tracked until the tillering stage, Zadocks 29-33. Water stress was initiated from the first day of imaging and the WL treatment pots reached the target field capacity within 16-24 days with an average of 19 days after initiation of imaging. The daytime Photosynthetic Active Radiation (PAR) was supplemented with LED red/blue light lamps, with an intensity of 200 μmol m^−2^ s^−1^. The experiment was conducted at the Nebraska Innovation Campus greenhouse, high-throughput plant phenotyping core facility (Scanalyzer 3D, LemnaTec Gmbh, Aachen, Germany), University of Nebraska-Lincoln.

A two-way factorial complete randomized experimental design, with 47 ILs and the recurrent parent, Svevo, was conducted. There were two water treatments: well-watered (control, WW) at 80% field capacity (FC) and water-limited (WL) at 30% FC (Supplementary Fig. S3), with three replicates for each combination. As quality control, we used empty pots, placed randomly in every second row. In total there were 296 pots. Plants were imaged daily for 35 days with visible Red, Green, and Blue (RGB) camera (Basler, Ahrensburg, Germany) taking 5 side-views (rotating 72°) and a single top-view. The image size was 2454×2056 pixels. After imaging, each pot was automatically weighed and watered to meet its calculated target weight. Greenhouse temperature kept at 22/16°C (day/night) during the experiment.

Based on the results of the first experiment, we selected two ILs (IL20, IL46) for detailed physiological characterization, alongside their parental line Svevo. A two-way factorial complete random design was conducted, with three genotypes, and two water treatments as described above, with four replicates for a total of 24 pots. The imaging started 7 DAT and imaging continued for 14 days.

### Image processing

PhenoImage GUI software (http://vis.unl.edu/~yu/Research.htm) was used for image processing based on MATLAB (The Mathworks, Inc., Massachusetts, USA). The workflow consisted of three main steps: image cropping, plant segmentation, and attribute extraction. In brief, image cropping was used to remove the frame of the chamber, followed by a background removal step based on color differences. Plant segmentation was based on filtration of pixel intensity (i.e., distinguishing between plant and non-plant pixels). As a result, the software can give the plant dimension, pixel sum, image moment, and convex area. All raw RGB images were deposited in the CyVerse and can be accessed at https://rb.gy/zvbief. Image data were stored in the following data structure: pot number_genotype_water treatment_replicate_date/side view_degree/image.png.

### Morpho-physiological trait characterization

We extracted some of the key morphological traits derived from RGB images included, PSA, plant height and width, plant architecture (convex area), plant density, and water-use efficiency (WUE) at the final day of the experiment. *Plant height* and *plant width* were calculated from plant dimensions. *Plant architecture* (convex area) was calculated to predict plant architecture trajectory. *Density* was calculated based on the ratio between pixel sum and plant architecture. *Plant biomass* was calculated based on the projected shoot area (PSA) as described by (Campbell et al., 2015). On the last day of the experiment, a subset of 19 ILs harvested, oven-dried (80° C), and weighed to obtain shoot dry weight. Correlation analysis showed a high correlation between PSA and shoot dry weight (r=0.96; *P*<10^−4^; Supplementary Fig. S14). The *relative growth rate* (RGR) was calculated by dividing daily pixel accumulation with pixel numbers from the previous day. Daily *water-use efficiency* (WUEt) was calculated as described by Momen et al. (2019), where (t) represents the day.

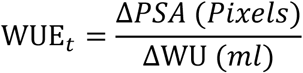

where ΔPSA is the daily PSA:

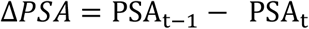

and ΔWU is the daily water used:

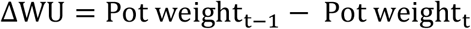

*Photosynthetic rate*, *transpiration rate,* and *stomatal conductance* were measured between 10 and 22 DAT (Zadocks 15-19) using a portable infra-red gas analyzer (LI-6800XT; Li-Cor Inc., Lincoln, NE, USA). Measurements were conducted at the mid-portion of the last fully expanded leaf from 9:00 to 13:00 (*n*=3).

*Root biomass* was measured at 22 DAT from 2L pots (45×19.5 cm) filled with 1.2 kg of Fafard germination soil (Sungro, Massachusetts, USA). Root tissue was harvested (*n*=4), washed and oven-dried (80° C) for 72h, and weighed to obtain root dry weight. *The root-to-shoot ratio* was calculated by dividing root dry weight with PSA (shoot dry weight).

### Characterization of root and shoot length

Uniform seeds were germinated in a Petri dish on moist germination paper for 5d in the dark at 22-25°C. Five seedlings of each genotype were placed on moist germination paper (25 × 38 cm; Anchor Paper Co., St. Paul, MN, USA), about 5 cm apart, with the germ end facing down. The paper was covered with another sheet of moist germination paper and rolled to a final diameter of 3 cm. The bases of the rolls were placed on a 4L beaker in a darkened growth chamber at a temperature of 24C/16C, 15h/9h day/night, at 50-60% relative humidity. A two-way factorial design was used with two genotypes (Svevo and IL20) and two water availabilities: WW and WL, with 8 replicate for each combination (total of 32). Eight cigar rolls were placed in a container (4 L) were the well water treatment was refiled on daily basis to keep the availability of 100 ml of water. The water-limited treatment filled once with 20 ml and did not re-filled during the experiment, or 20 ml (without refilling) for WL. Each container was wrapped with plastic to prevent water evaporation. Shoot and root length were measured daily by scale, from 3 to 8 DAT (Zadocks 11).

### Statistical Analyses

The JMP^®^ ver. 15 statistical package (SAS Institute, Cary, NC, USA) was used for statistical analyses unless otherwise specified. The longitudinal response was fitted for genotypes (collectively or separately) under each water treatment. Analysis of Variance (ANOVA) was used to assess the possible effects of genotype (G), environment (E), and G×E interactions on morpho-physiological traits of genotypes. Frequency distribution was determined for all morpho-physiological traits on the last day. Components of descriptive statistics are graphically presented in box plot: median value (horizontal short line), quartile range (25 and 75%) and data range (vertical long line). Principle Component Analysis (PCA) was used to determine associations between traits. PCA was based on a correlation matrix and is presented as biplot ordinations of the ILs (PC scores). Three components were extracted using eigenvalues >1.2 to ensure the meaningful implementation of the data by each factor. An agglomerative hierarchical procedure with an incremental sum of squares grouping strategy was employed using Ward’s method (Ward, 1963), for classification. Pearson correlation for all morpho-physiological traits was conducted for each water treatment. Drought-susceptibility index (S) was calculated according to Fischer and Maurer (1978):

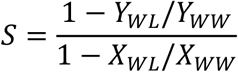

where *Y*_*WL*_ and *Y*_*WW*_ are the mean phenotypic values of a certain genotype under the respective treatments, and *X*_*WL*_ and *X*_WW_ are the mean performances of all genotypes. Morpho-physiological correlation matrix and Density distribution were plotted with R software (RStudio Team, 2015).

### Broad-sense heritability dynamics

Broad-sense heritability (_*bs*_*h*^2^) and its components, genetic component 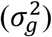, and G×E interaction 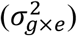, were calculated for each day of imaging across the two water treatments using ANOVA-based variance components:

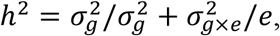

where 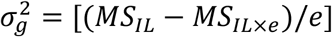, 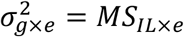, *e* is the number of water treatments and MS is the mean square.

### RNA extraction and sequencing

Root tissues were collected daily from 8 to 11 days after germination (Zadocks 11) and frozen in liquid nitrogen until RNA extraction. RNA was extracted using the plant/fungi total RNA purification kit (Norgen Biotek Corp., Canada) with on-column DNase treatment (Qiagen, Germany). Sample contamination and RNA integrity were assessed using the Nan D-1000 spectrophotometer (Thermo Fisher Scientific). Based on the physiological analysis, we selected samples from day six for RNAseq, with two repeats for each combination (total 8). Single-end (150bp) bar-coded cDNA libraries were prepared for sequencing on the Illumina HiSeq2000 sequencer (NGS Core, Nebraska Medical Center Omaha, USA).

### Accession Number

Raw sequencing files of mRNA sequencing are available at the short read archive of the National Center for Biotechnology Information (https://www.ncbi.nlm.nih.gov) under accession number GSE163450.

### Data processing and analysis

FastQ quality of each sample was manually inspected using FastQC (http://www.bioinformatics.babraham.ac.uk/projects/fastqc). Barcode removal, filtering, and trimming of low-quality reads were executed using the command line tools Trimmomatic (Bolger et al., 2014). Each RNA-seq read was trimmed to make sure the average quality score exceeded 30 and has a minimum length of 70bp. Sequences were aligned to the available Svevo and Zavitan reference genomes (Avni et al., 2017; Maccaferri et al., 2019). Using TopHat (Trapnell et al., 2009), allowing for up to 2 bp mismatches per read. Reads mapped to multiple genomic locations were removed. Numbers of reads per gene were counted by the software tool of HTSeq-count using corresponding rice gene annotations and the “union” resolution mode was used (http://www-huber.embl.de/users/anders/HTSeq). Differential expression analysis of count data and data visualization were conducted with the DESeq2 package (Love et al., 2014). To detect significant DEGs, a 5% false discovery rate (FDR) correction for multiple comparisons was determined (Benjamini and Hochberg, 1995), and a minimal |0.5| log2FC threshold was applied. Venn diagrams were created with http://bioinformatics.psb.ugent.be/webtools/Venn. Gene ontology, Singular Enrichment Analysis (SEA), and Parametric Analysis of DEGs set Enrichment for biological processes and pathways was conducted with AgriGO (http://systemsbiology.cau.edu.cn/agriGOv2; Tian et al., 2017).

### Gene ontology network

Biological processes and molecular function networks were established using the DEGs GO terms with REVIGO software (http://revigo.irb.hr); this summarizes lists of GO terms using a clustering algorithm that relies on semantic similarity measures (Supek et al., 2011). The analysis outputs were transferred to the Cytoscape software (https://cytoscape.org), which served as a network biology analysis and visualization tool (Otasek et al., 2019).

### Genetic analysis of candidate DEGs

Candidate genes were analyzed on the wheat efp browser for expression in different tissues and phenological stages (http://bar.utoronto.ca/efp_wheat/cgi-bin/efpWeb.cgi; Ramírez-González et al., 2018). Gene sequences were compared with the publically available genome of Svevo https://wheat.pw.usda.gov/GG3/genome_browser and compared to Zavitan gene sequences with a blast against the Zavitan genome https://wheat.pw.usda.gov/cgi-bin/seqserve/blast_wheat.cgi. Differences in splice variance number of candidate genes were perceived from the blast on the GrainGenes website https://wheat.pw.usda.gov/cgi-bin/seqserve/blast_wheat.cgi. DNA translation to amino acids was done with the free online software https://web.expasy.org/translate

## Supplemental Data

The following supplemental materials are available.

**Supplemental Table S1.** Correlations between morpho-physiological traits under well-watered and water-limited treatments.

**Supplemental Table S2.** Longitudinal coefficient of variance for PSA.

**Supplemental Table S3.** Comparison of PSA, plant architecture and plant architecture density under two water treatments for each cluster.

**Supplemental Table S4.** Regression equation of relative growth rate.

**Supplemental Table S5.** Comparisons of A, T, and *gsw* between Svevo, IL20, and IL46 under two water treatments throughout the experiment.

**Supplemental Table S6.** Comparisons of morpho-physiological traits between Svevo, IL20, and IL46 under two water treatments.

**Supplemental Table S7.** Physical location of wild emmer introgressions of IL20 on the Zavitan genome.

**Supplemental Table S8.** Gene annotation within IL20 introgressions.

**Supplemental Table S9.** Significant differentially expressed genes.

**Supplemental Table S10.** Root-related candidate genes.

**Supplemental Table S11.** List of ILs and their chromosomal introgressions.

**Supplemental Figure S1.** Correlation between projected shoot area (PSA) and shoot dry weight.

**Supplemental Figure S2.** Longitudinal dynamics of morpho-physiological traits under contrasting water treatment.

**Supplemental Figure S3.** Experimental design.

**Supplemental Figure S4.** Plant projected shoot area (PSA) dynamics of introgression lines and Svevo under well-watered and water-limited treatments.

**Supplemental Figure S5.** Correlation matrix between morpho-physiological traits under well-watered and water-limited treatments.

**Supplemental Figure S6.** Principal component analysis of morpho-physiological traits.

**Supplemental Figure S7.** Hierarchical clustering of morpho-physiological traits under water-limited and in terms of susceptibility index and clusters expression pattern.

**Supplemental Figure S8.** Longitudinal dynamic of plant architecture and density.

**Supplemental Figure S9.** Longitudinal heritability of plant density and architecture.

**Supplemental Figure S10.** Longitudinal dynamics of Svevo, IL20 and IL46 assimilation rate under well-watered and water-limited treatments.

**Supplemental Figure S11.** Longitudinal dynamics for stomatal conductance and transpiration rate under well-watered and water-limited treatments.

**Supplemental Figure S12.** Longitudinal dynamics of the root-to-shoot ratio under contrasting water treatment.

**Supplemental Figure S13.** Heat map of candidate genes from Zavitan expression atlas and read count of the candidate gene TRIDC2AG073520 at different developmental stages.

**Supplemental Figure S14.** Expression atlas of TRIDC2AG073520 in the wheat efp browser.

## Acknowledgments

We thank members of the Peleg and Walia labs for technical assistance with experiments. We thank Dr. Paul Staswick for critical reading of the manuscript. This research was partially supported by the Chief Scientist of the Israel Ministry of Agriculture and Rural Development to ZP, the U.S. Agency for International Development Middle East Research and Cooperation (grant # M34-037) to ZP, and the Agricultural Research Division Wheat Innovation Fund from the University of Nebraska-Lincoln to HW and TA.

