## Supplementary material for "Wild emmer introgressions alter root-to-shoot growth dynamics in response to water stress": SI Figures

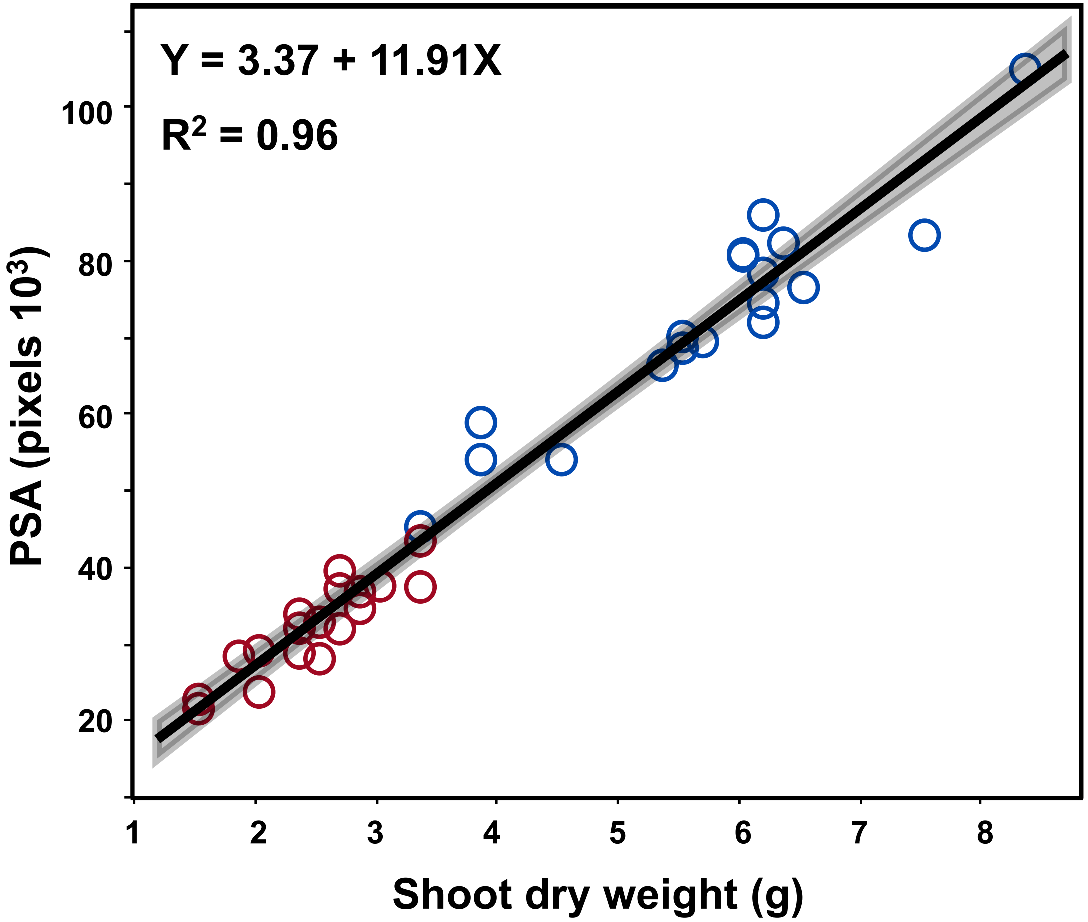


**Supplementary Figure S1** Correlation between projected shoot area (PSA) and shoot dry weight of 18 wild emmer introgression lines under well-watered (blue) and water-limited (red) treatments.

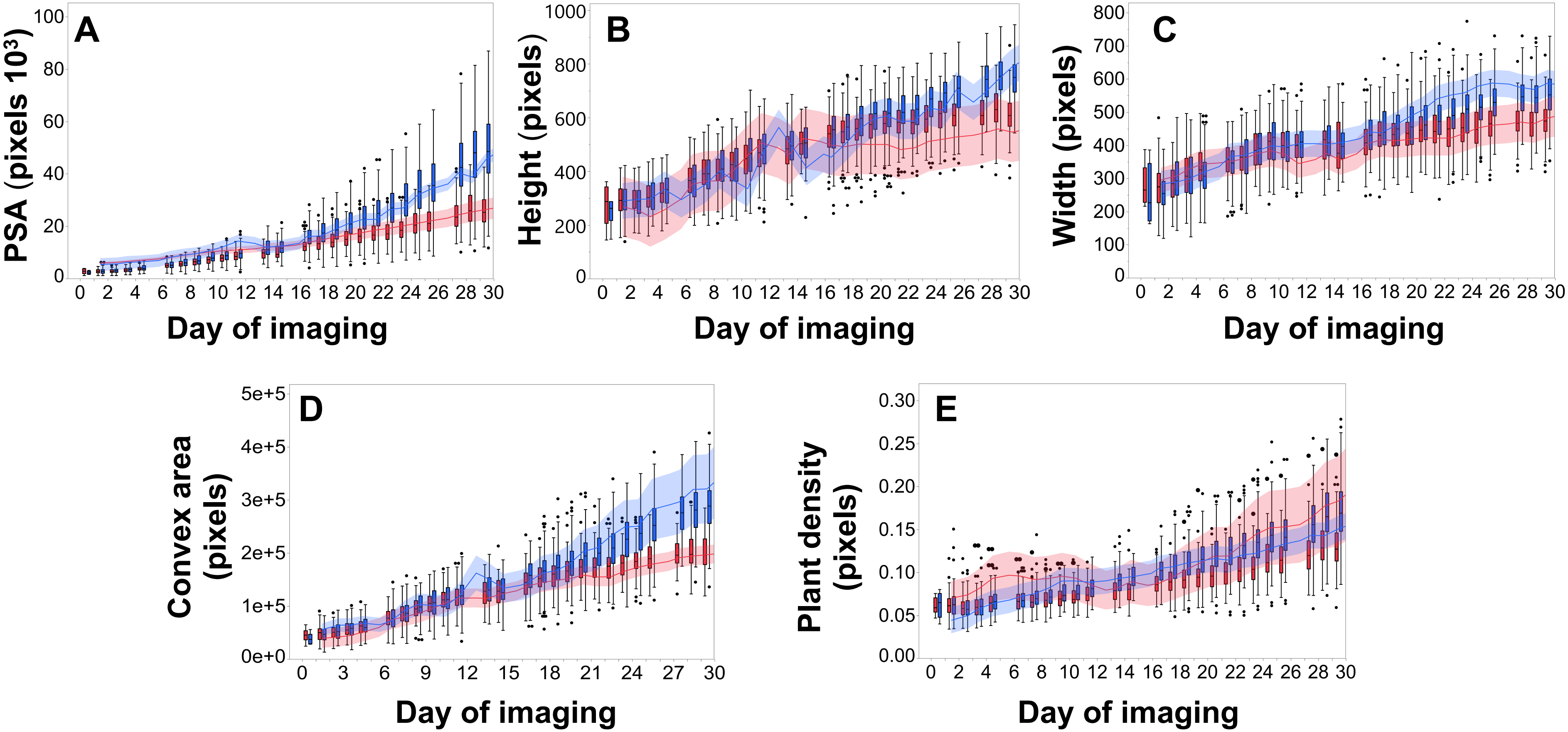


**Supplemental Figure S2**. Morpho-physiological dynamics of all introgression lines (in box plot) and Svevo (continuous line) under well-watered (WW; blue) and water-limited (WL; red) treatments. Plant projected shoot area (PSA), Plant height, Plant width, Plant architecture (Convex area), and Plant density The parental line Svevo (Sv) are marked by arrows. The box plots for each morpho-physiological trait represents the lower quantile (25%) median (50%) and upper quantile (75%) of the IL population within water treatment. The dashed line represents the first day of significance (*P*≤0.05) between the water treatments.

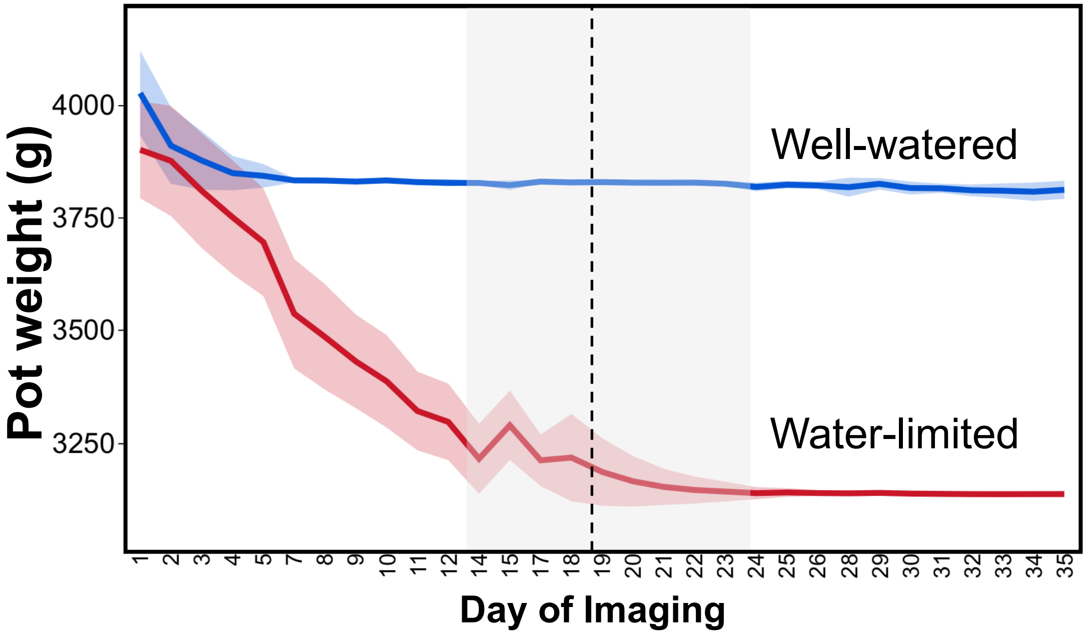


**Supplementary Figure S3.** Experimental design. Plants were grown under well watered conditions (80% field capacity, FC) for six days and than water stress was applied by withholding water till 30% FC (day 23) and kept at 30% for 12 additional days. Dashed line represent the average day of reaching to 30% field capacity. Grey area represent the variation among ILs at the first 30% field capacity point.

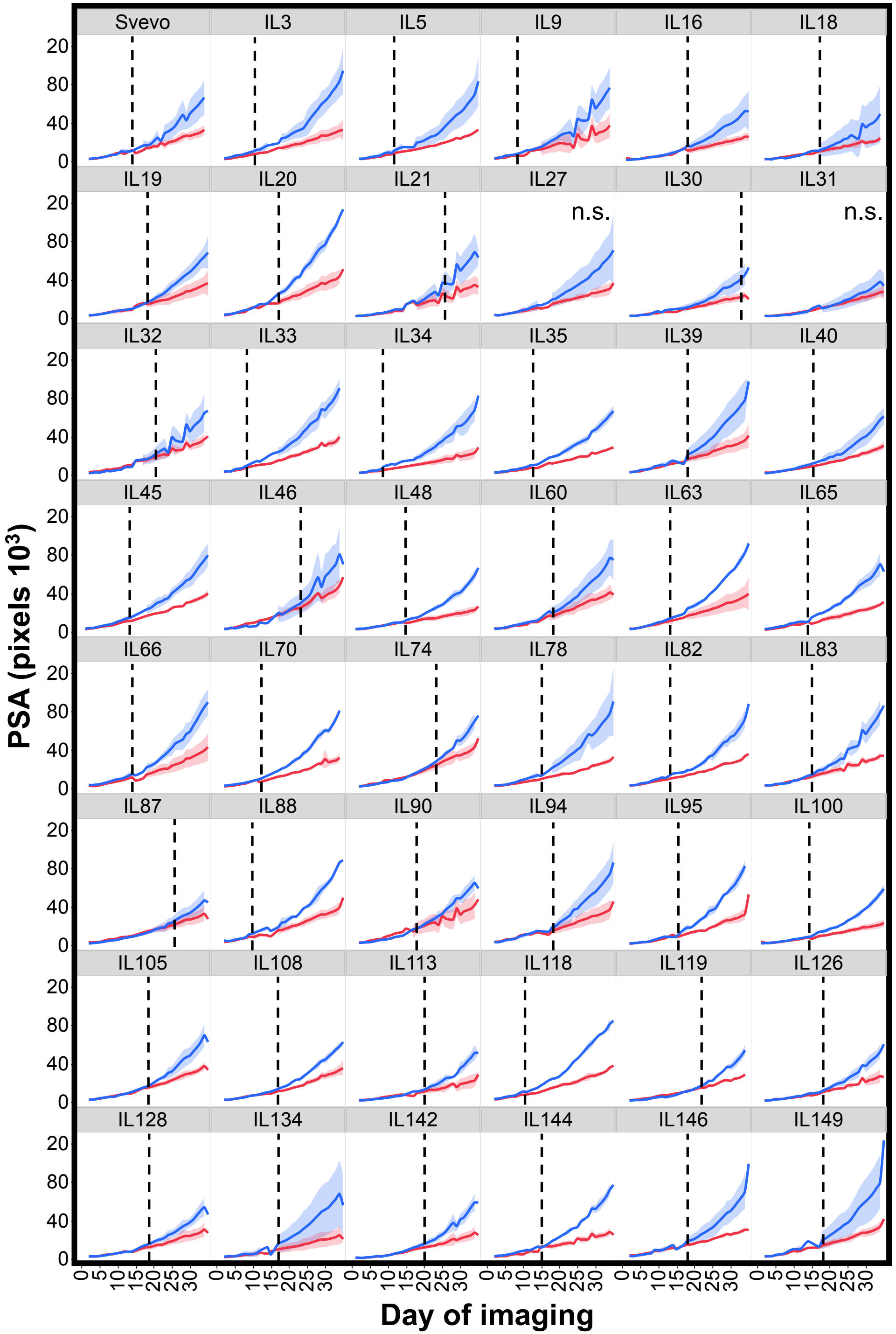


**Supplemental Figure S4**. Plant projected shoot area (PSA) dynamics of introgression lines (IL) and Svevo under well-watered (WW, blue) and water-limited (WL, red) treatments (*n*=3). Continues lines represent the smooth curve under WW and WL through the data and the shaded area represents the standard error of the smooth curve. The dashed line represents the first day of growth differentiation between water treatments, n.s. represents not significant.

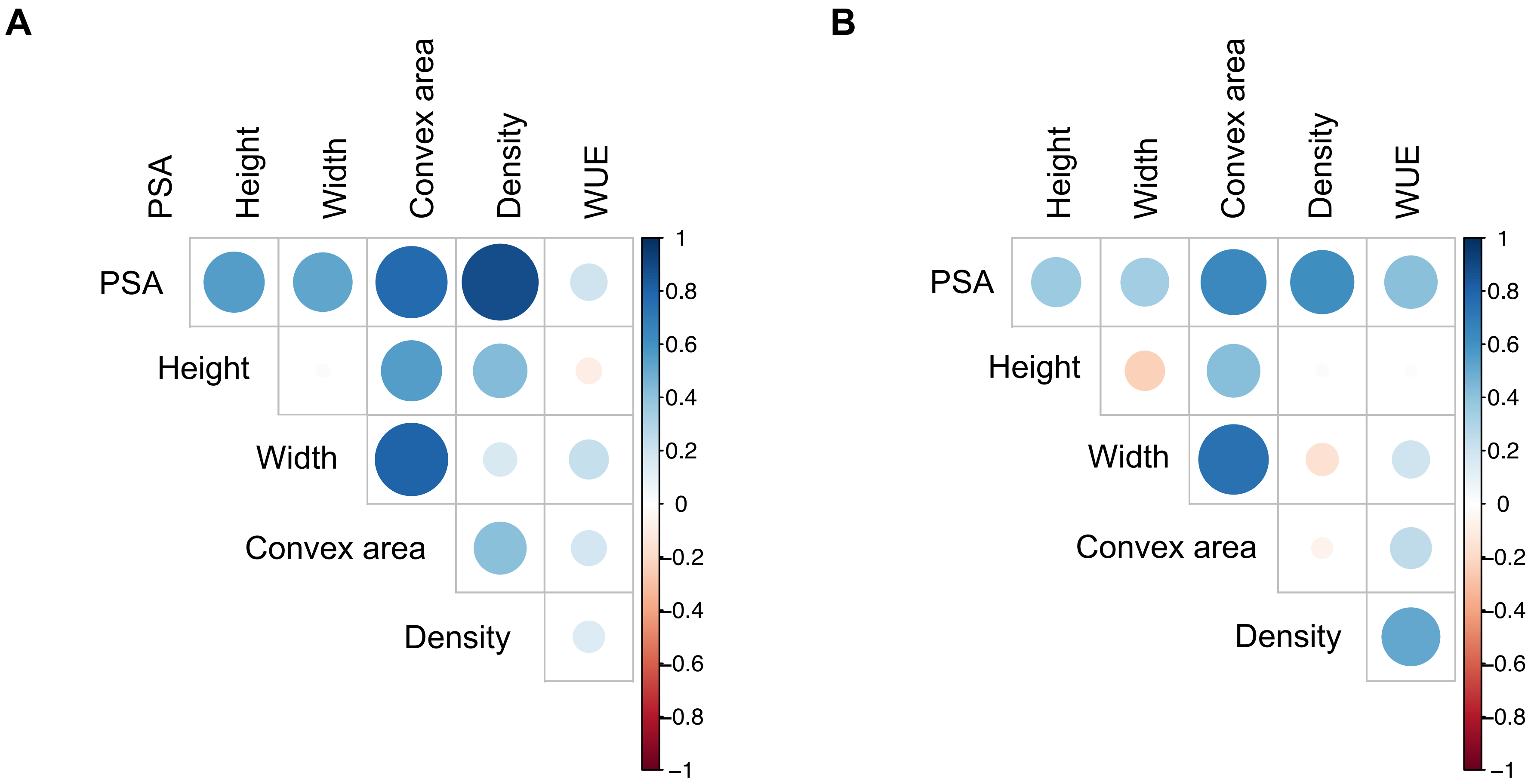


**Supplemental Figure S5**. Pearson correlation matrix between morpho-physiological traits under (**A**) well-watered and (**B**) water-limited. Projected shoot area (PSA), Plant height (Height), Plant width (Width), Plant architecture (convex area), Plant density (density), Water-use efficiency (WUE). Colors indicate level of correlation (r) from positive correlation (red) to negative (blue). The size of the circle indicate level of significance.

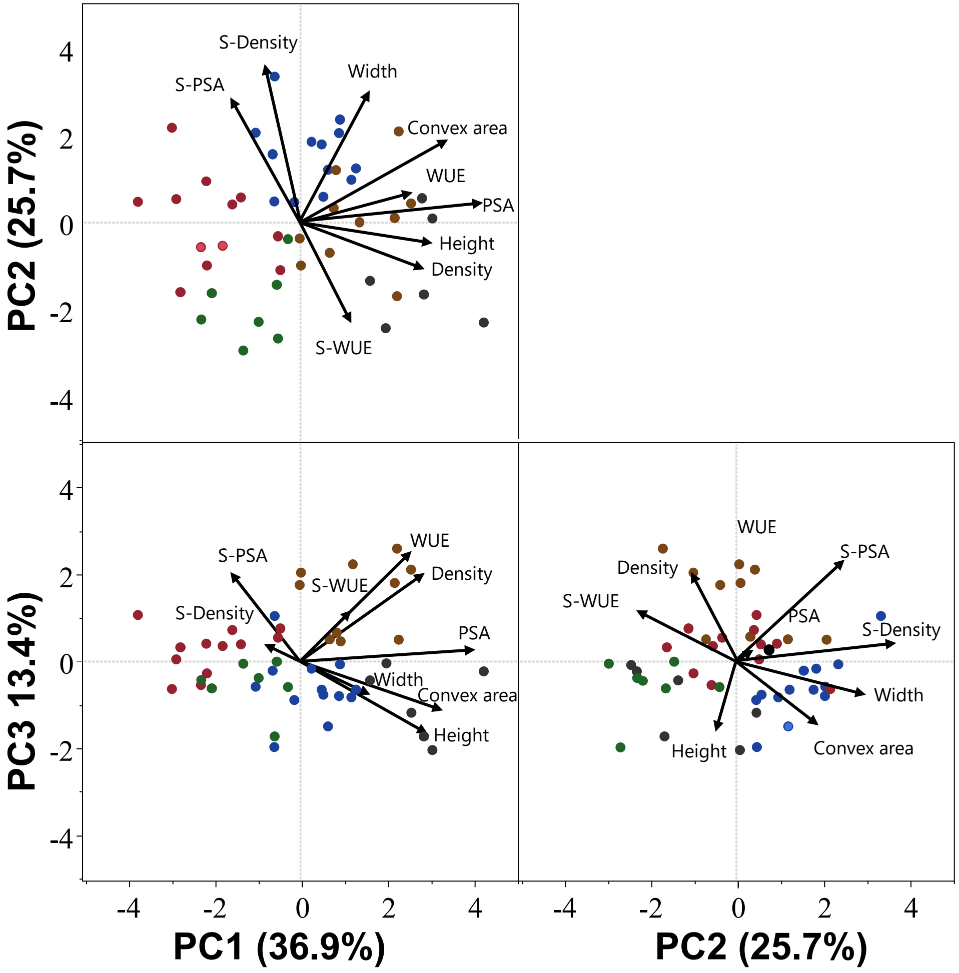


**Supplemental Figure S6.** Principal component (PC) analysis of continuance morpho-physiological traits (eigenvalues >1.2) under water-limited conditions and as expressed by drought susceptibility index (S). Biplot vectors are trait factor loadings for PC1, PC2 and PC3. Projected shoot area (PSA), Plant height (Height), Plant width (Width), Plant architecture (Convex area), Plant density (Density), water-use efficiency (WUE). The five clusters of stress responsiveness: low productivity - high stability (LPHS; Green), low productivity - moderate plasticity (LPMP; Red), high productivity - moderate plasticity (HPMP; Blue), high productivity - high plasticity (HPHP; Orange), high productivity - high stability (HPHS; Black).

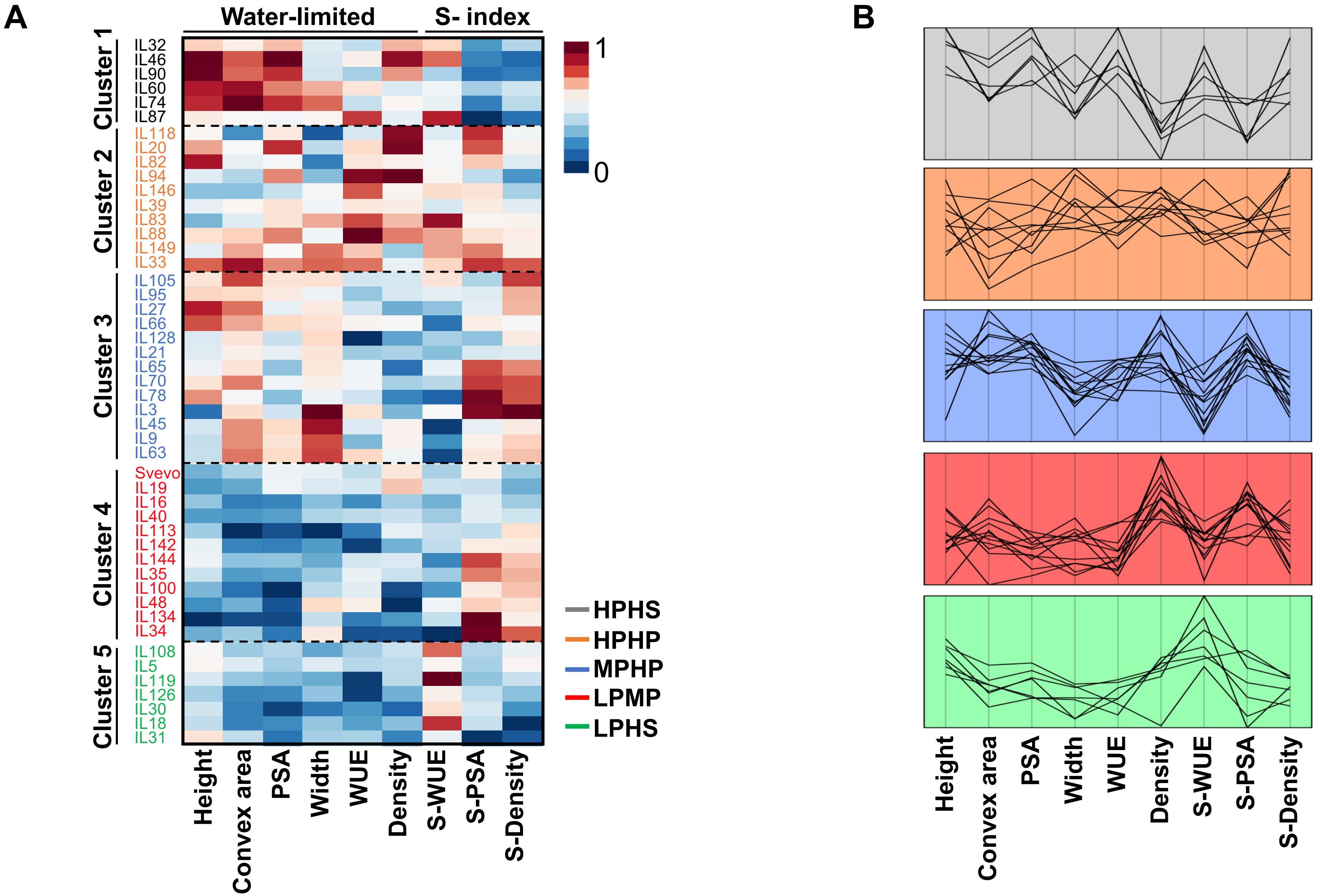


**Supplemental Figure S7.** (**A**) Hierarchical clustering integrated heat map of morpho-physiological traits under water-limited treatment and as expressed in drought susceptibility index (S) at the last day of the experiment (*n*=3). Projected shoot area (PSA), Plant height (Height), Plant width (Width), Plant architecture (Convex area), Plant density (Density), water-use efficiency (WUE). ILs were clustered to five clusters according to ward method and named by the cluster characteristics. The heat map colors from blue to red represent lower to higher scaled values respectively. (**B**) The mean expression pattern of each IL by its cluster.

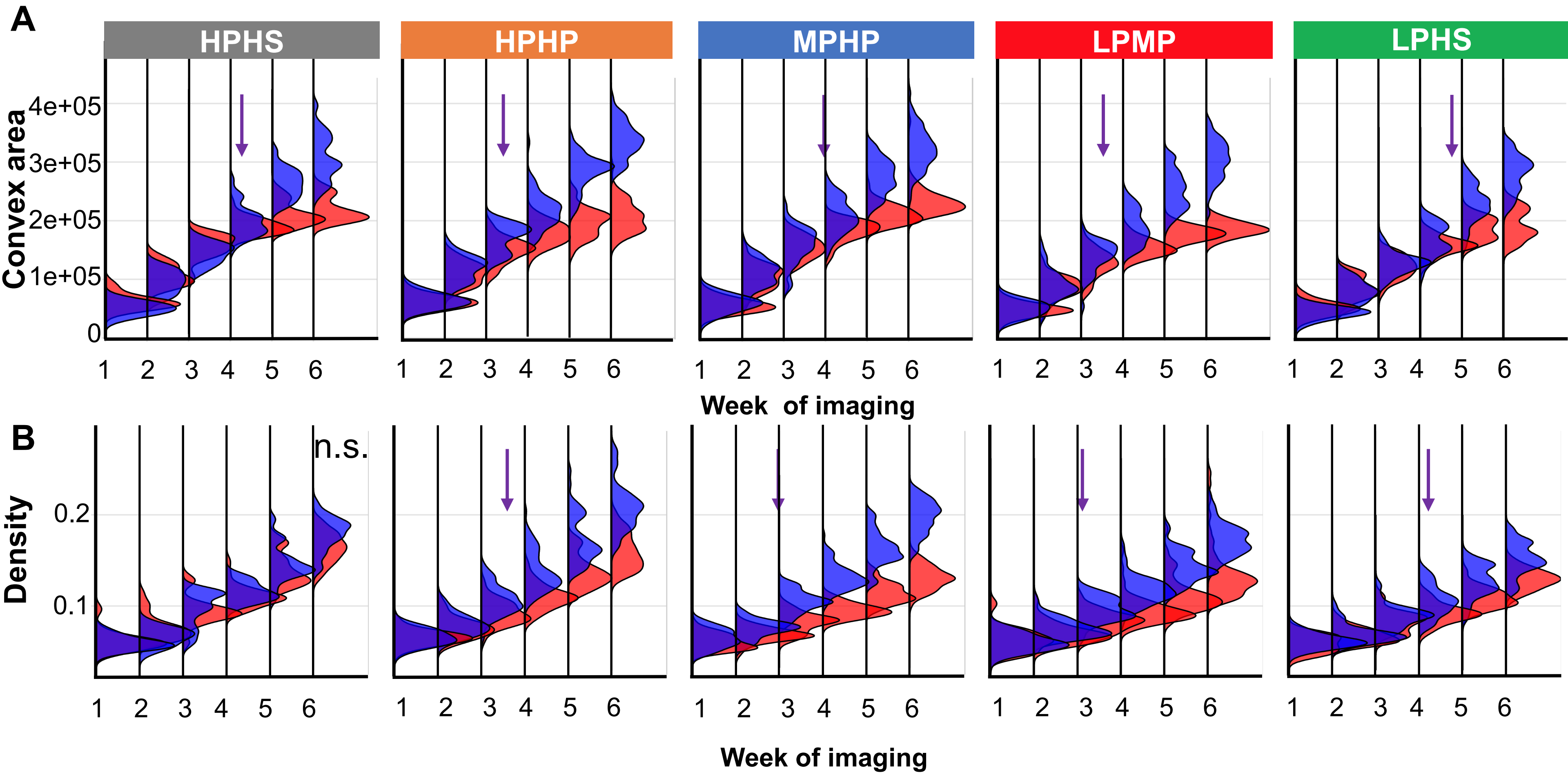


**Supplemental Figure S8.** Longitudinal dynamics of the five responsiveness clusters. (**A**) Longitudinal frequency distribution of plant architecture (Convex area), and **(B)** Plant architecture density (Density), of each responsiveness cluster under well-watered (WW; blue) and water-limited (WL; red) treatments. The five clusters of stress responsiveness: high productivity - high stability (HPHS; gray), high productivity - high plasticity (HPHP; Orange), moderate productivity - high plasticity (MPHP; Blue), low productivity - moderate plasticity (LPMP; Red), low productivity - high stability (LPHS; Green). The point of significant ($P \leq 0.05$) response to water stress is marked above with arrow, or non-significant (n.s.).

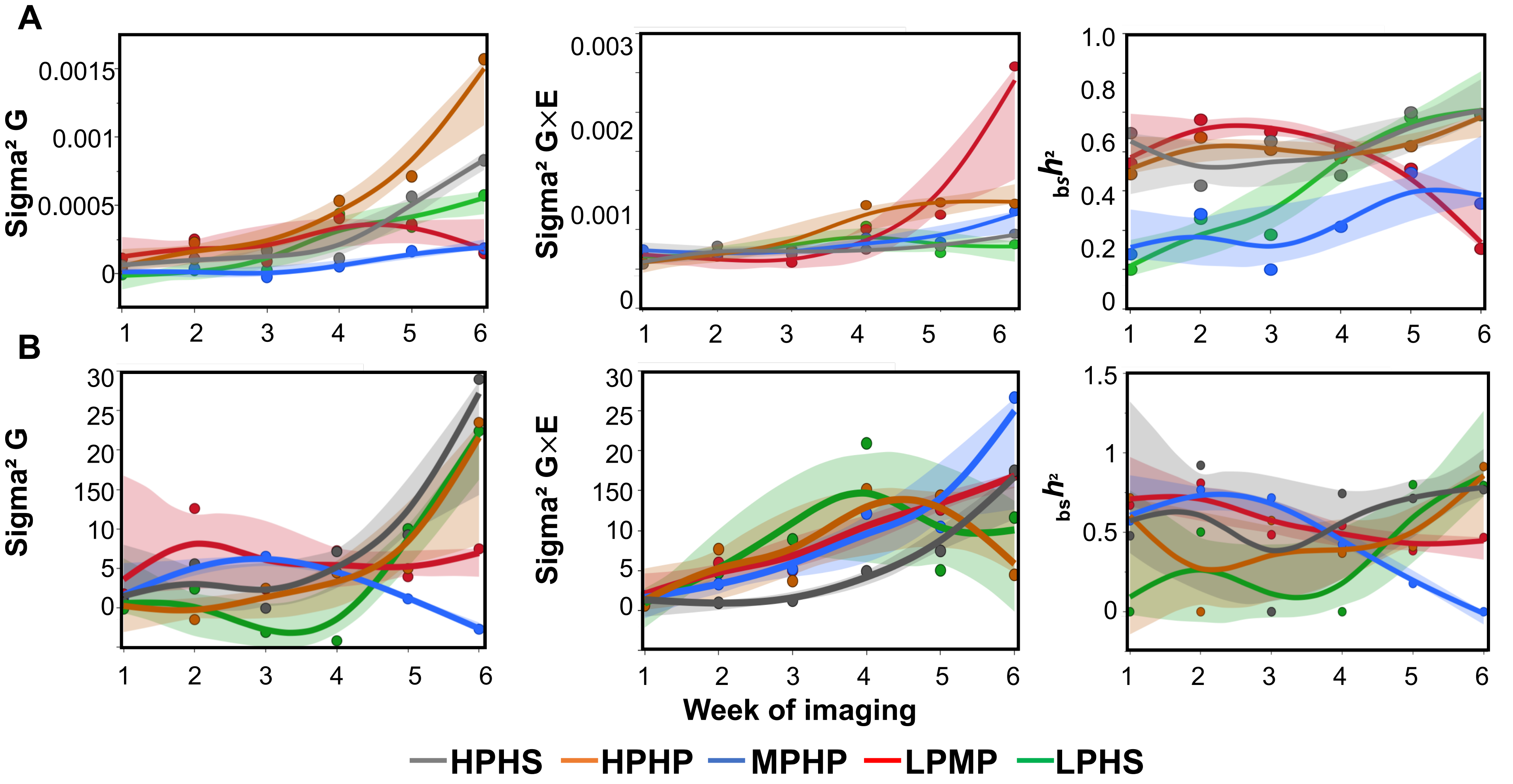


**Supplemental Figure S9**. Longitudinal heritability components of genetic component (Sigma² G), environmental interaction (Sigma²G × E), and broad sense heritability ($_{bs}h^2$) for (**A**) Plant density, and (**B**) Plant architecture. The five clusters of stress responsiveness: high productivity - high stability (HPHS; gray), high productivity - high plasticity (HPHP; Orange), moderate productivity - high plasticity (MPHP; Blue), low productivity - moderate plasticity (LPMP; Red), low productivity - high stability (LPHS; Green).

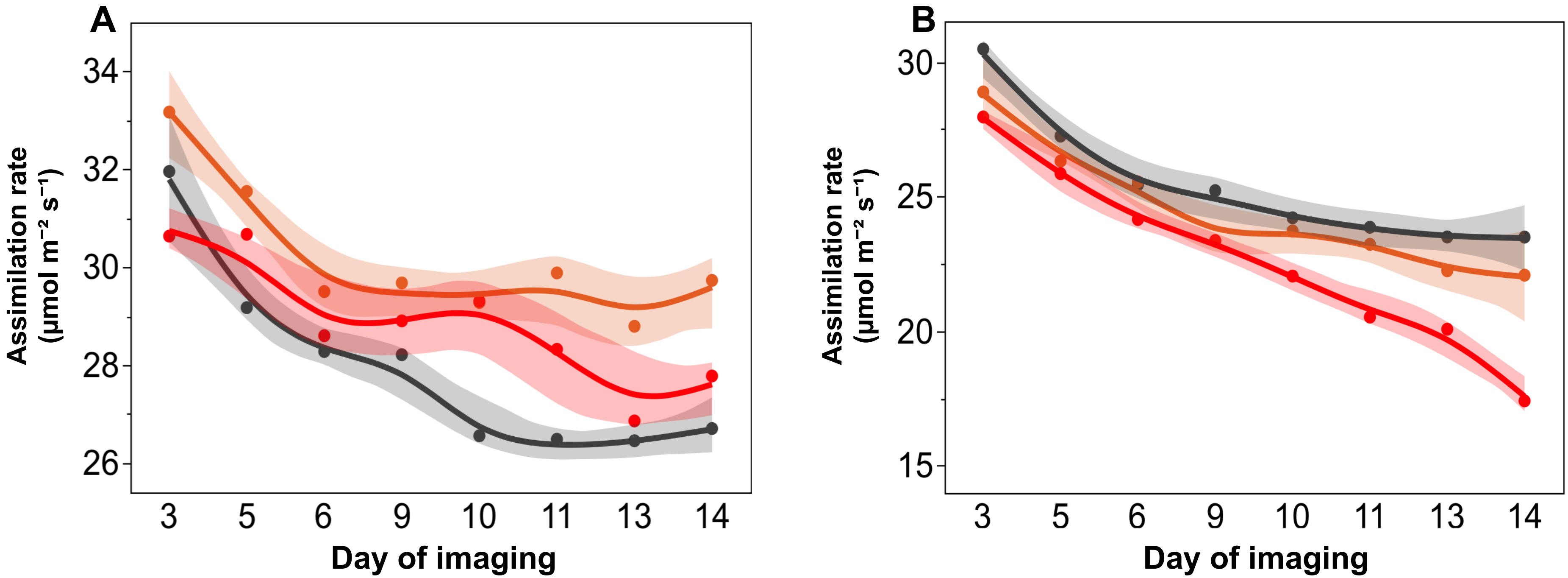


**Supplemental Figure S10.** Longitudinal dynamics of Svevo (red), IL20 (orange) and IL46 (grey) assimilation rate (**A**) under well-watered (WW) and (**B**) water-limited (WL) treatments. Markers represents the genotypic mean under specific water treatment (*n*=3). Continues line represent the smooth curve through the data and the shaded area represents the standard error of the smooth curve.

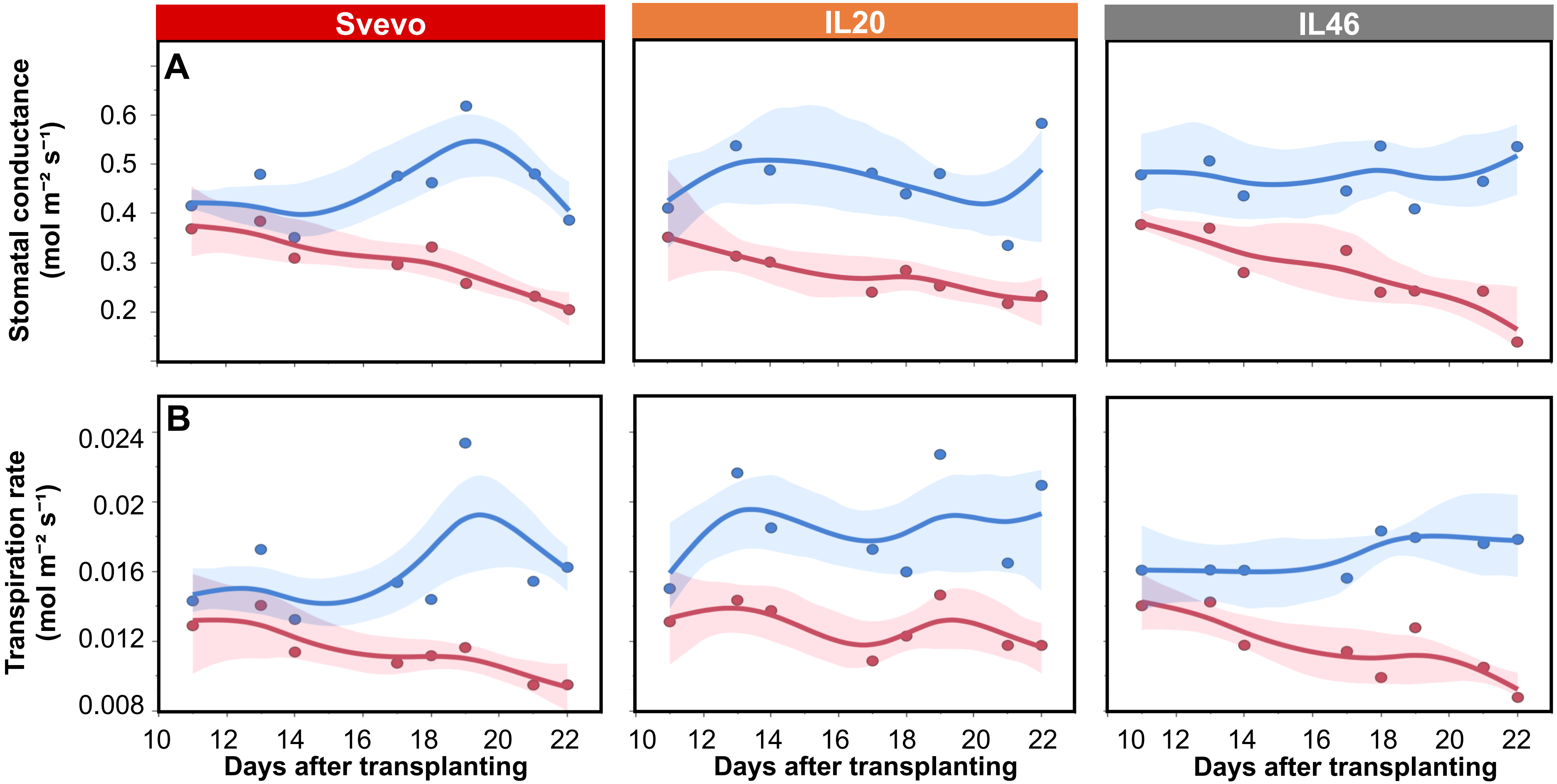


**Supplemental Figure S11.** Longitudinal dynamics of Svevo, IL20 and IL46 for (**A**) Stomatal conductance, (**B**) transpiration rate under well-watered (WW; blue) and water-limited (WL; red) treatments. Dashed lines represents the fitted linear growth of each genotype under specific water treatment. Markers represents the genotypic mean under specific water treatment (*n*=4). Continues line represent the smooth curve through the data and the shaded area represents the standard error of the smooth curve .

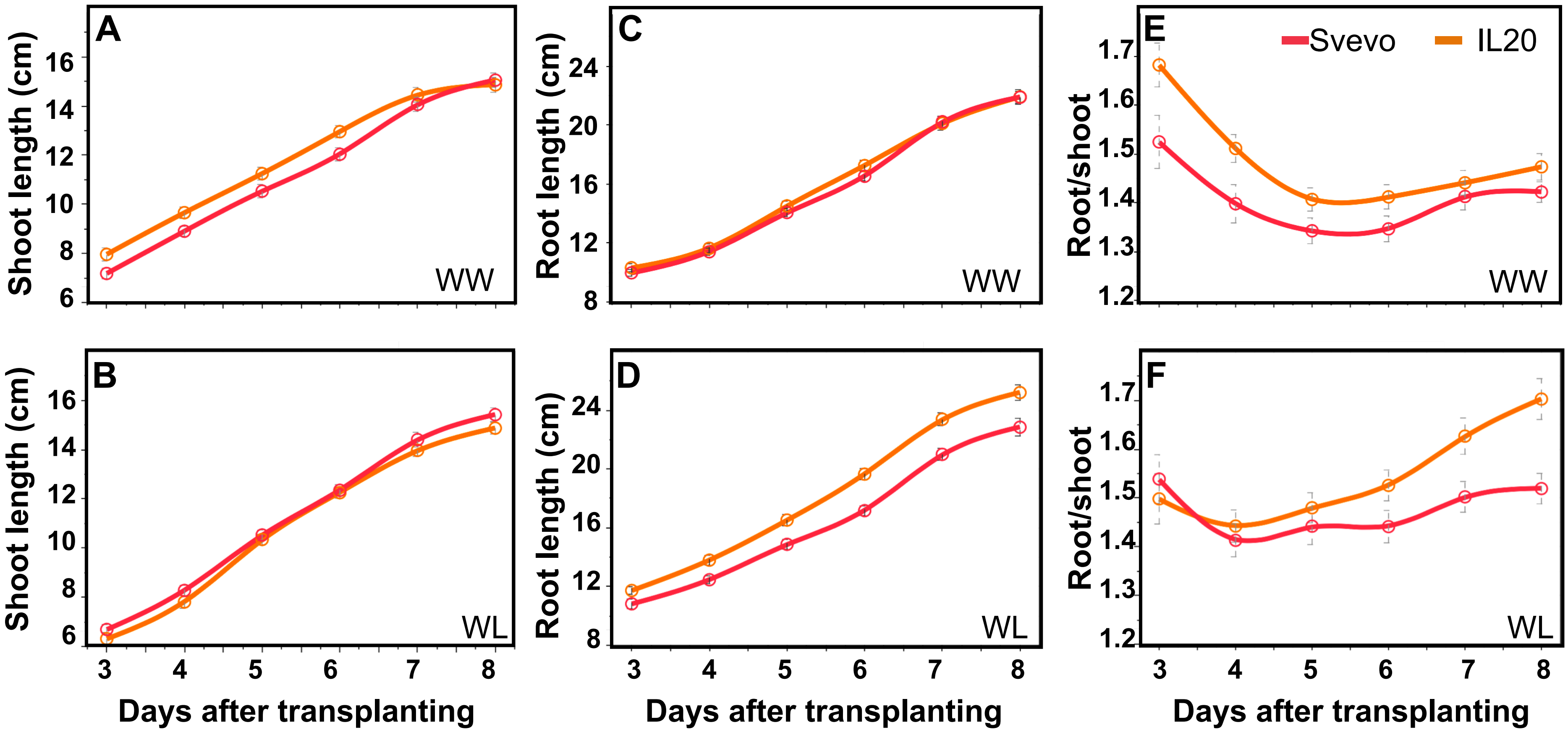


**Supplemental Figure S12.** Longitudinal dynamics of Svevo and IL20 (red and orange, respectively) response to water-stress. (**A**) shoot length under WW, (**B)** shoot length under WL, (**C**) root length under WW, (**D**) root length under WL, (**E**) root-to-shoot ratio (Root/shoot) under WW, (**F**) root-to-shoot ratio (Root/shoot) under WL.

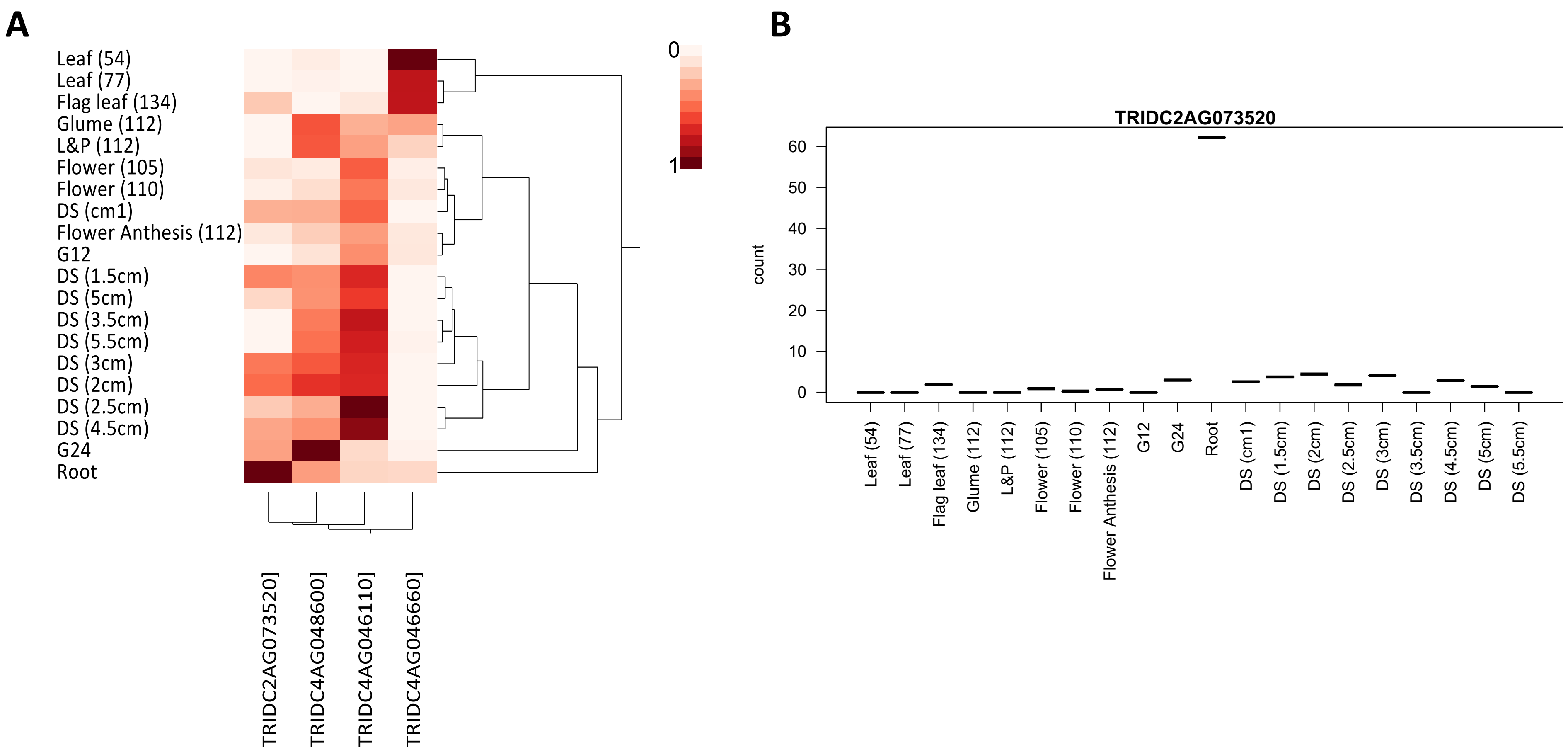


**Figure S13.** (**A**) Hierarchical clustering integrated heat map of four high confidence candidate genes from Zavitan wild emmer expression atlas under different tissues, morphological stages and days from planting )in the brackets): Leaf (54), Leaf (77), Flag leaf (134), Glume (112) , Lemma and Palea (L&P) (112), Flower (105), Flower (110), Flower Anthesis (112), grain 12 days after anthesis (G12), grain 24 days after anthesis (G24), Root - seedlings at 11 days after germination, developing spike (DS) (1 cm1), DS (1.5 cm), DS (2 cm), DS (2.5 cm), DS (3 cm), DS (3.5 cm), DS (4.5 cm), DS (5 cm), DS (5.5 cm). (**B**) Read count of the candidate gene TRIDC2AG073520 at different developmental stages.

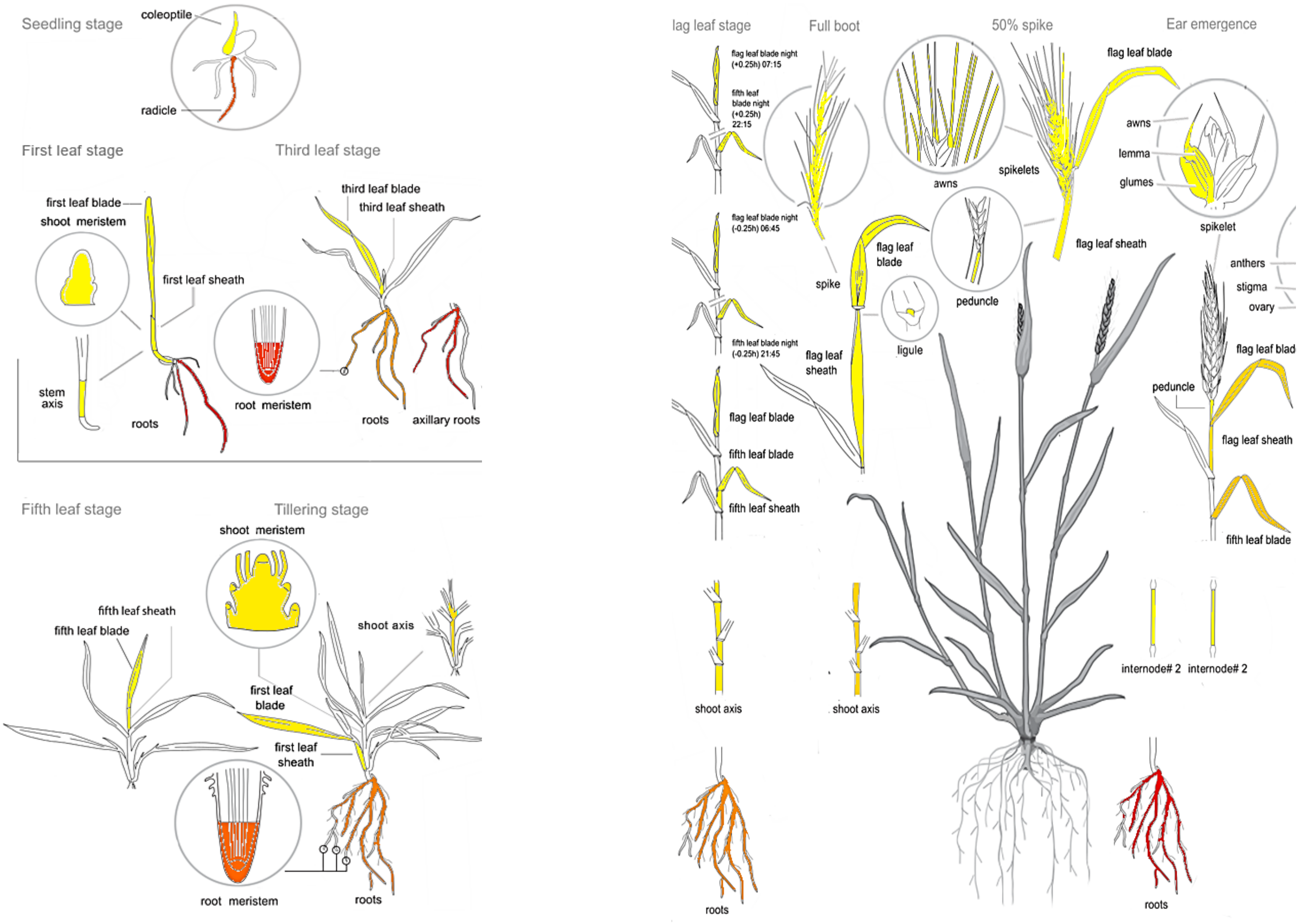


**Supplemental Figure S14.** Expression atlas of TRIDC2AG073520 gene in the wheat efp browser.
